# A phosphorylation-deficient ribosomal protein eS6 is largely functional in *Arabidopsis thaliana*, rescuing mutant defects from global translation and gene expression to photosynthesis and growth

**DOI:** 10.1101/2023.05.30.542942

**Authors:** Anwesha Dasgupta, Ricardo A Urquidi Camacho, Ramya Enganti, Sung Ki Cho, Lindsey L. Tucker, John S. Torreverde, Paul E. Abraham, Albrecht G. von Arnim

## Abstract

The eukaryote-specific ribosomal protein of the small subunit eS6 is phosphorylated through the Target of rapamycin (TOR) kinase pathway. Although this phosphorylation event responds dynamically to environmental conditions and has been studied for over 50 years, its biochemical and physiological significance remains controversial and poorly understood. Here we report data from *Arabidopsis thaliana*, which indicate that plants expressing only a largely phospho-deficient isoform of eS6 grow essentially normally under laboratory conditions. The eS6A (*RPS6A*) paralog of eS6 functionally rescued double mutations in both *rps6a* and *rps6b* genes when expressed at approximately twice the wild-type dosage. A mutant isoform of eS6A lacking the major six phosphorylatable serine and threonine residues in its carboxyl-terminal tail also rescued the lethality, rosette growth, and polyribosome loading of the double mutant. It also complemented many mutant phenotypes of *rps6* that were newly characterized here, including photosynthetic efficiency, and the vast majority of gene expression defects that were measured by transcriptomics and proteomics. However, compared to plants rescued with a phospho-enabled version of eS6A, the phospho-deficient seedlings retained a mild pointed-leaf phenotype, root growth was reduced, and certain cell cycle related mRNAs and ribosome biogenesis proteins were misexpressed. The residual defects of the phospho-deficient seedlings could be understood as an incomplete rescue of the *rps6* mutant defects, with little or no evidence for gain-of-function defects. As expected, the phospho-deficient eS6A also rescued the *rps6a* and *rps6b* single mutants; however, phosphorylation of the eS6B paralog remained lower than predicted, further underscoring that plants can tolerate phospho-deficiency of eS6 well. Our data also yield new insights into how plants cope with mutations in essential, duplicated ribosomal protein isoforms.

## INTRODUCTION

The central function of ribosomes is conserved across the kingdoms of life. However, eukaryotic ribosomes contain expansion segments in their ribosomal RNAs, eukaryote-specific ribosomal proteins, and covalent modifications that are not found in prokaryotic ribosomes. Ribosomal protein of the small subunit 6 (eS6 or RPS6) is a pan-eukaryotic protein and was the first ribosomal protein reported to be phosphorylated more than four decades ago (Kabat, 1970; Gressner and Wool, 1974). Its amino-terminus is buried in the 40S-60S subunit interface while its carboxy-terminal tail harboring a string of phosphorylatable serine and threonine residues at the end of a long alpha-helix is located on the solvent exposed side of the 40S. The presence of phosphorylatable residues is highly conserved in all eukaryotes even though the number of the residues varies among species.

Phosphorylation of eS6 occurs in a polysome context (Duncan and McConkey, 1982). Phosphorylation is also highly dynamic. In animals, eS6 phosphorylation is sensitive to nutrients, hormones, growth factors, and a variety of stress conditions (Meyuhas, 2008). In plants, initial evidence for phosphorylation of eS6 came from tomato plants and maize root tips, where phosphorylation was suppressed by abiotic stresses, heat shock and hypoxia, respectively (Scharf and Nover, 1982; Bailey-Serres and Freeling, 1990). In Arabidopsis, where up to seven sites can be phosphorylated, eS6 phosphorylation levels are induced in response to both internal and external signals: light, circadian clock, sucrose (Turkina et al., 2011; Dobrenel et al., 2016; Chen et al., 2018; Enganti et al., 2018), elevated CO_2_ (Boex-Fontvieille et al., 2013), auxin (Schepetilnikov et al., 2013), and cytokinin (Yakovleva and Kulaeva, 1987). eS6-P has evolved to integrate signals in an elaborate manner. As a case in point, phosphorylation is induced in the daylight and repressed during a dark night, but in contrast is induced by the circadian clock during subjective night and repressed during subjective day (Choudhary et al., 2015; Enganti et al., 2018), an incoherent signaling network that may allow eS6-P to sense subtle shifts in photoperiod or diel illumination (Panchy et al., 2020).

eS6 phosphorylation in plants is considered a canonical readout of the Target-of-rapamycin - S6 kinase (TOR-S6K) pathway because this is the only established pathway known to regulate eS6 phosphorylation in plants (Mahfouz et al., 2006; Dobrenel et al., 2016; Chen et al., 2018). Despite detailed insights into the regulation of eS6-phosphorylation by upstream signals and kinases, the biochemical and physiological role of eS6-P has remained fairly enigmatic. Knock-in mice expressing only the alanine-substituted, nonphosphorylatable version of eS6 exhibit severe whole-body phenotypes, including a reduced size, glucose intolerance, and muscle weakness, along with a higher rate of protein synthesis (Ruvinsky et al., 2005; Ruvinsky et al., 2009). The phosphorylation of eS6 is implicated in cell size control, hyperplasia in pancreatic cancer, glucose homeostasis, and activation of neurons (Ruvinsky et al., 2005; Ruvinsky et al., 2009; Knight et al., 2012; Khalaileh et al., 2013; Wittenberg et al., 2016). The consequences of eS6-P at the biochemical level including translation are not well understood. Mouse embryonic fibroblasts lacking phosphorylatable eS6 had decreased translation fidelity, an increased rate of translation overall, and proliferated faster (Wittenberg et al., 2016). The eS6-P was slightly more abundant on shorter coding sequences, and eS6-P slightly boosted their translation efficiency (Bohlen et al., 2021). eS6 phosphorylation has also been linked to transcriptional regulation of genes encoding ribosome biogenesis factors (Chauvin et al., 2014).

In budding yeast, a detailed study of eS6-P deficiency including ribosome footprinting did not discover any effect of the phosphorylation potential on bulk translation nor a role in regulation of gene expression via (transcription or) translation. An effect on ribosome biogenesis was attributed to reduced protein expression (Yerlikaya et al., 2016). Recently, Arabidopsis eS6-phosphorylation was proposed to support translation re-initiation, based on differences between phospho-null and phospho-mimic versions with respect to (i) their in-vitro interaction with the re-initiation supporting protein, RISP and (ii) their ability to support expression of a reinitiation-dependent reporter mRNA (Mancera-Martinez et al., 2021).

Here, we report our findings from an extensive characterization study of plants lacking either one or both wild-type *RPS6* paralogs and harboring either wild-type or phospho-deficient eS6 transgenes. In Arabidopsis the genes encoding eS6, *RPS6A* and *RPS6B*, are functionally largely equivalent but non-redundant (Creff et al., 2010). We find that a phospho-deficient allele of eS6 is able to rescue the lethality of plants lacking both wild-type *RPS6* genes. We also find it to be largely functional in complementing the growth defects in single-paralog mutants. Double *rps6a rps6b* mutants complemented with a P-deficient eS6 had reverted to normal photosynthetic efficiency (Qy_max_) and had largely normal mRNA and protein levels for photosynthesis proteins. These P-deficient plants also had no striking defects in global translation, again complementing defects seen in the single *rps6* mutants. However, transcriptome analysis revealed that the plants displayed subtle defects in gene expression of several cytokinesis related genes. They also tended to have asymmetric cotyledons and transiently pointed first leaves, defects hinting at incomplete complementation of the *rps6* mutations and potentially indicating a whole-plant defect stemming from the P-deficiency. These results mirror and extend findings on the function of this core phosphorylation event from yeast and vertebrates to photosynthetic organisms.

## RESULTS

### A new allele for *rps6a* recapitulates the typical phenotypes of a previous null allele

We sought out a new null allele for the A paralog of eS6 (protein: eS6A; gene: *RPS6A*) in order to facilitate double mutant construction later on. The new null-allele of *RPS6A*, *rps6a-2* was marked by sulfadiazine resistance rather than kanamycin resistance **(Supplemental Fig. 1A, B, C)** and did not express detectable mRNA **(Supplemental Fig. 1D)**. The *rps6a-2* allele recapitulated the phenotype of an earlier allele, *rps6a-1 (Creff et al., 2010)* with respect to pointed primary leaves, root length, and seed set **(Supplemental Fig. 2A-E).**

Because we were planning to test the function of eS6 phosphorylation using eS6A as a stand-in for both paralogs, we wanted to confirm whether the functions of eS6A and eS6B are equivalent and interchangeable, as suggested by Creff et al., 2010. Indeed, the new *rps6a-2* allele was fully complemented by either the *RPS6A* or *RPS6B* gene **(Supplemental Fig. 3A)**. However, the *rps6b-1* allele was largely but not fully complemented by either *RPS6A* or *RPS6B* **(Supplemental Fig. 3B)**. In these experiments the transgenes consisted of the native promoter and the native exon-intron structure. Taken together these results confirm that eS6A and eS6B are functionally equivalent. We surmise that the complementation in the *rps6b-1* background was incomplete because the residual truncated transcripts made from the *RPS6B* gene **(Supplemental Fig. 1D)** may interfere with gene expression of the transgene. This was not analyzed further. From this point forward, we will refer to the *rps6a-2* allele just as *rps6a* and *rps6b-1* as *rps6b*.

### Phosphorylation-deficient alleles of eS6A and eS6B and their subcellular localization

We generated an extensive series of site-directed mutant alleles for both *RPS6A* and *RPS6B*. To prevent phosphorylation, serine or threonine residues in the C-terminal tails of eS6A and eS6B (**Fig. 1A**) were changed to alanine (ι1S>A alleles). In an effort to create phospho-mimic alleles, S or T were changed to aspartate (ι1S>D alleles; **Supplemental Fig. 1E**).

**Figure 1.**
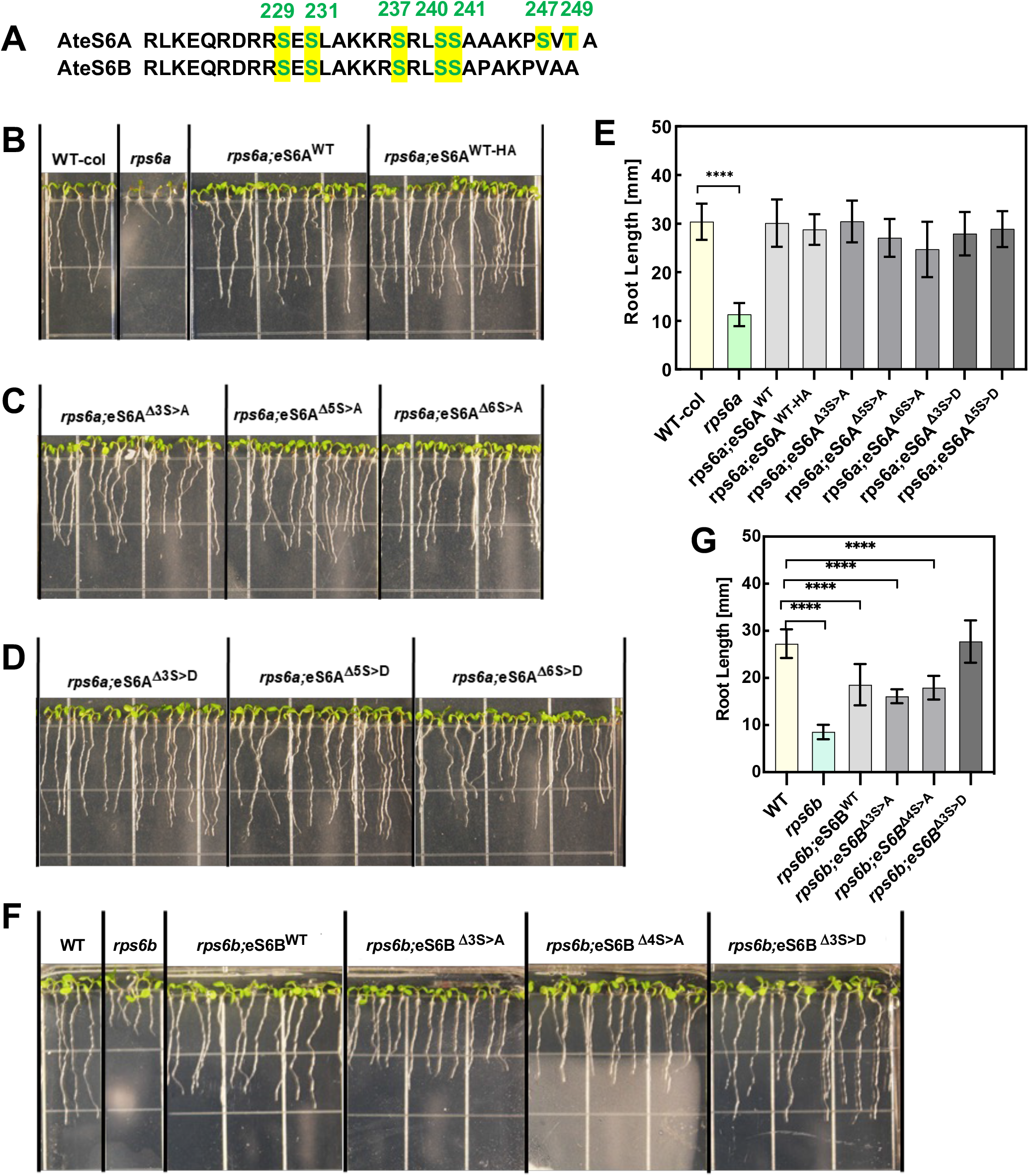
Phosphodeficient alleles of eS6A and eS6B substantially complement *rps6* mutant root growth defects. **(A)** Phosphorylatable serines and one threonine in the C-terminal tails of eS6A and eS6B. **(B)** Wild-type, **(C)** phosphodeficient and **(D)** phosphomimic alleles of eS6A complement the *rps6a* mutation. **(E)** Quantitation of root lengths from lines in (B-D). **(F, G)** Phosphodeficient and phosphomimic alleles of eS6B partially complement the *rps6b* mutation.

Expressed in *Nicotiana benthamiana* as an enhanced YFP (EYFP) fusion, both the wild-type and a fully phosphodeficient allele, eS6A ^1′7S>A^, accumulated strongly in the nucleus and especially in the nucleolus, as well as in the cytosol, as expected for ribosomal proteins **(Supplemental Fig. 4A)**. The RPS6A wild-type and eS6A ^1′7S>A^ mutant proteins also appeared in structures resembling mini-nucleoli and other more or less numerous small granules, collectively referred to as nuclear punctae. eS6A ^1′7S>A^ formed punctae more often than eS6A wild type **(Supplemental Fig. 4B)**. eS6A ^1′7S>A^ transformed cells also tended to have more than one nucleolus but this trend was not statistically significant **(Supplemental Fig. 4C)**. eS6 was tagged at the N-terminus because tagging the C-terminus was more likely to disrupt the phosphorylation at that site. If EYFP-eS6 were incorporated into the 40S, then the bulky YFP may disrupt subunit joining with the 60S. Taken together, the phosphorylation status did not appear to affect the subcellular targeting of eS6 in a major way.

### Phosphodeficient alleles of eS6A and eS6B substantially complement the growth phenotypes of single *rps6* mutants

If the phosphorylation of eS6 were essential for each of the two paralogs, then abolishing the potential for phosphorylation in one paralog should result in a deficiency in growth or development when P-deficient constructs in the native genomic context (promoter, introns, UTRs) are transformed into the single-mutant backgrounds. We measured root length as a proxy for functionality of the transgenes, because root length is a quantitative trait and highly sensitive to the dosage of eS6. The *rps6a* root growth defect was complemented well by several P-deficient and P-mimic isoforms of eS6A (**Fig. 1B-E**).

The *rps6b* mutation was also complemented, but root growth was slower than wild type with three eS6B ^1′S>A^ alleles tested (**Fig. 1F-G**). The incomplete complementation cannot be attributed to the phosphodeficiency of eS6B because wild-type *RPS6B* genes did not fully complement the *rps6b* mutation either, as shown earlier **(Supplemental Fig. 3C)**. These data contradict the hypothesis that each paralog of eS6 must be phosphorylatable for full function. Phosphorylation is a physiologically dispensable property of eS6 under our growth conditions.

### *rps6a* and *rps6b* mutant seedlings have defects in the translation apparatus

In order to later discern subtle effects of the phosphodeficiencies in eS6 on the translation apparatus, we thoroughly analyzed the translation apparatus of the *rps6a* and *rps6b* strains by polysome profiling with sucrose density gradient centrifugation **(Supplemental Fig. 5A-C)**. Profiles were collected both 0.5 h before and 2.5h after the daily dark to light shift (i.e. at zeitgeber times ZT23.5 and ZT2.5) because light exposure induces a large and rapid increase in mRNA ribosome loading (Liu et al., 2012). To bring out the full scale of this shift seedlings were grown in the absence of sucrose for these experiments.

We detected an increase in the relative abundance of the 60S subunit, which had been suggested earlier (Creff et al., 2010) and was statistically significant for both *rps6a* and *rps6b* in the light **(Supplemental Fig. 5C)**. Polysomes were decreased slightly in both *rps6a* and *rps6b*, which was statistically significant for the small polysome fraction (SP) after lights-on. In response to the dark-to-light shift, the fold-increase in polysomes (P/(NP+P)) was 1.53 or greater for both *rps6* mutants **(Supplemental Fig. 6A-B**), showing that both *rps6* mutants responded as robustly as WT to the dark-to-light shift.

From these data we surmise that production of sufficient 40S subunit is rate limiting in the *rps6* single mutants, because only a single paralog is available for eS6, not only for growth but also for ribosome production. Meanwhile, the 60S subunit overaccumulates; i.e., the amount of 40S and 60S subunit are slightly out of balance in both *rps6* mutants as compared to wild type.

eS6 was phosphorylated normally in the *rps6a* and *rps6b* mutants. In detail, at the end of night, the phosphorylation of S237 and S240 was low overall and almost absent in polysomes. It increased dramatically, overall as well as in polysomes, with the dark-to-light shift **(Supplemental Fig. 5A-B**). These results match those in wild type (Enganti et al., 2018) and show that both eS6A and eS6B are phosphorylated under light in a polysome context and that both isoforms are recognized by both phospho-specific antibodies.

### Phosphorylation of eS6 is reduced to an unexpected degree in partially phosphodeficient seedlings

In the single *rps6* mutants that were complemented with a phospho-deficient allele, the second paralog of eS6 remains available to be phosphorylated. We performed immunoblots on whole cell extracts, and also on gradient-fractionated material, to address whether eS6B remains phosphorylated when phosphorylation of eS6A is disrupted.

In whole cell extract at ZT2.5, three different eS6A^1′S>A^ mutants had unexpectedly low levels of P-eS6B (**Figure 2A**). This was not because the antibodies detected P-eS6B less well, as evident in the single-mutant *rps6a* lane (also see **Supplemental Fig. 5A**). In eS6A^1′S>D^ mutants, phosphorylation was reduced only mildly and less consistently. eS6B phosphorylation was also reduced in polysome gradients from these lines (**Fig. 2B**) with data from many additional blots summarized in **Fig. 2C**. Although we also examined phosphorylation of eS6A protein when *rps6b* mutants were complemented with phosphodeficient eS6B, the results were not as extensive and did not show a convincing loss of phosphorylation in eS6A (**Fig. 2C**).

**Figure 2.**
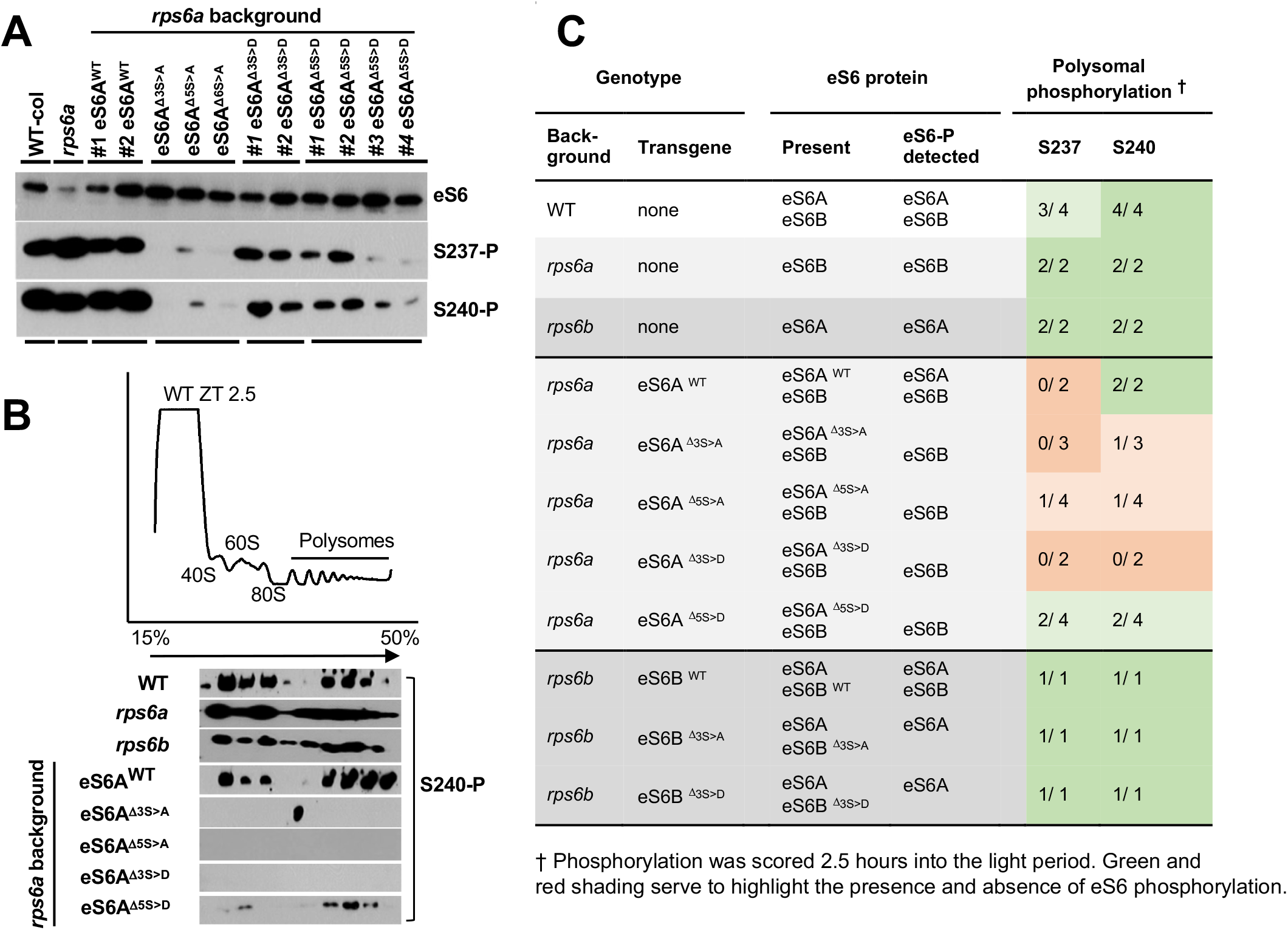
Phosphorylation status of eS6 in *rps6* single mutants that were complemented with wild-type or phosphodeficient alleles of eS6. Both S>A and S>D mutations are shown. **(A)** Whole cell extracts from seedlings in the rps6a background were probed for phosphorylation of S237 and S240 on western blots using phospho-specific antibodies (Enganti et al., 2018). Several lines were analyzed in duplicate. **(B)** Polysome gradients from seedlings harvested at ZT2.5. The absorption profile at the top comes from *rps6a* harboring eS6A ^Δ6S>A^ and is shown as a representative. The other gradients looked similar. Shown below are immunoblots of *rps6a* single mutants complemented with various phospho-deficient alleles; gradient fractions 1-12 were probed for phospho-S240. **(C)** Phosphorylation of eS6 in polysomes. Summary of data from western blots such as those in panel (B). The fractional data (e.g. 1/ 4) indicate the proportion of replicate experiments in which phosphorylation of eS6 was detected. In the remaining replicates, phosphorylation was undetectable or weak.

Taken together, these data suggest that the P-deficiency in eS6A^1′S>A^ interferes with the phosphorylation of the eS6B paralog. Although we do not know the mechanism, this observation of poor eS6-P is important, because it implies that the *rps6a* plants that were complemented with phospho-deficient eS6A were substantially free of eS6-P, at least at the main sites S237 and S240, yet grew essentially normally (**Fig. 1C-E**).

### The phospho-deficiency of eS6A has minor or no effects on global polysome loading

We then examined using sucrose density gradients whether overall polysome loading was altered in the phosphodeficient single mutants. Quite clearly, in the phospho-deficient single mutants (*rps6a*; *RSP6B*; eS6A^1′3S>A or 1′5S>A^) the dark-to-light transition triggered the typical rise in polysome loading **(Supplemental Fig. 6A)**, just as in wild type and in the *rps6* mutants. Therefore, the P-deficient plants have no striking defects in polysome loading. The significant elevation in the level of the 60S subunit seen earlier in the *rps6* mutants was no longer significant with the phospho-enabled and phospho-deficient eS6B and eS6A transgenes **(Supplemental Fig. 6A)**. These results suggest that the phosphodeficient isoform rescues the ribosome biogenesis defects.

### Plants that are almost entirely deficient for eS6-P grow largely normally

Next, we generated double *rps6a rps6b* mutants that harbored transgenes with variably phospho-deficient versions of eS6A and eS6B. The majority of the work was done with an allele where 5 serines and 1 threonine were replaced with alanine (ι16S>A, **Supplemental Fig. 1E**). As controls, double mutants were complemented with an HA-tagged version of eS6A (eS6A ^WT-HA^) or with wild-type RPS6B and with a Δ3S>D allele of RPS6B. The eS6A^1′6S>A^ complemented plants were indeed deficient for eS6-P for both S237 and S240 (**Fig. 3A**).

**Figure 3.**
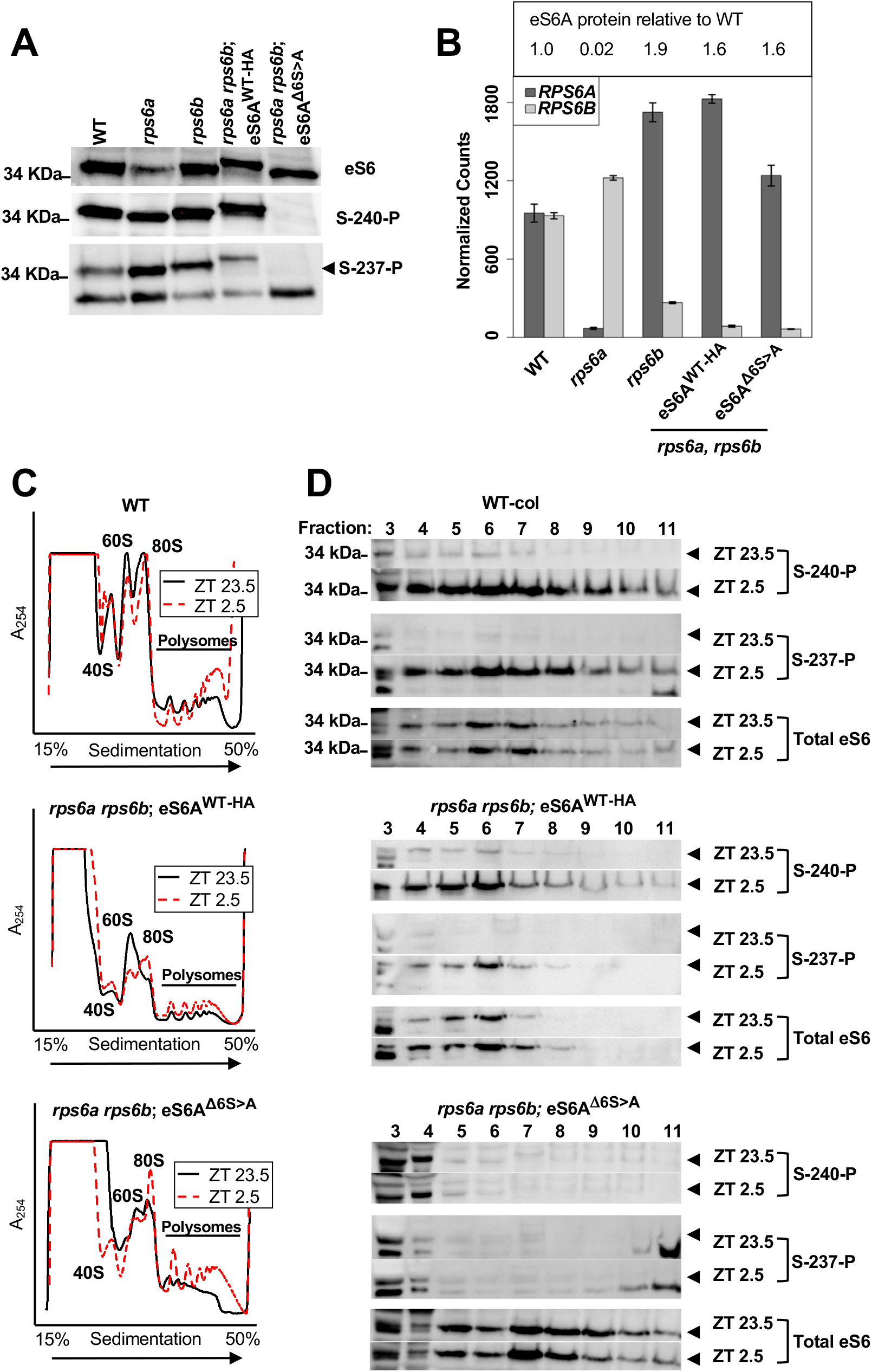
Biochemical characterization of plants lacking eS6 phosphorylation. **(A)** Phosphorylation of eS6 is not detected in *rps6a rps6b* double mutants that harbor the phospho-deficient eS6A^1′6S>A^ transgene. Whole protein extracts from seedlings were probed with phospho-specific antibody for P-S240 or P-S237. Note: Our S-237 antibody detects two bands consistently. Repeated results and experience with the eS6A^1′6S>A^ allele have convinced us that the upper band running at ∼35 KDa is the desired band. **(B)** mRNA read counts for eS6A and eS6B from an RNA-Seq experiment with the indicated genotypes. Proteomics data from the same genotypes expressed as a log_2_-fold difference to wild type. Note that the three transgenes provide elevated levels of eS6 protein, as expected given the elevated mRNA read counts.

**(C)** Polysome loading in *rps6a rps6b* double mutants complemented with phospho-enabled and phospho-deficient transgenes. Polysome loading was examined 30 minutes prior to lights-on (ZT23.5) and 2.5h after lights-on (ZT2.5). Representative polysome profiles show that eS6-P deficient plants are able to increase their ribosome loading during the dark-to-light shift.

**(D)** Immunoblots demonstrating that eS6 is phosphorylated in a polysome context in double *rps6a rps6b* mutants complemented by an HA-tagged eS6A (middle). In contrast, in the phosphodeficient plants (right) no phosphorylation is detectable at the position corresponding to eS6 (arrowhead). Note that fractions 3-4 at the top of the gradient contain several crossreacting proteins.

RNA-Seq and proteomics experiments to be described below demonstrated that the eS6A^1′6S>A^ mRNA and protein were expressed at an elevated level, equivalent to about 1.3- and 1.6-fold, respectively, of the native eS6A (**Fig. 3B**). Therefore, it appears that the homozygous eS6A^1′6S>A^ transgene provides about as much eS6 mRNA and protein as can be expected in a heterozygous *rps6b* mutant, which is considered a recessive mutation.

For Wt-col, polysomal eS6 phosphorylation at S240 and S237 increased robustly within 2.5h of light exposure. This is consistent with the increase in global polysome loading and higher eS6 phosphorylation in response to light. A similar pattern was observed for double mutant seedlings harboring HA-tagged WT-eS6A. Both of these results confirm that phosphorylated eS6 gets incorporated into actively translating polysomes. The eS6A^1′6S>A^ complemented seedlings lacked S240 and S237 phosphorylation, as expected (**Fig. 3D**). However, total eS6 can still be detected in polysomal fractions, suggesting that the eS6A^1′6S>A^ protein, despite lacking six major sites of phosphorylation, gets incorporated into polysomes (**Fig. 3D**).

Considering that *rps6a rps6b* double mutants are embryo-lethal (Creff et al., 2010), it was striking that the P-deficient plants were fully viable. However, a number of subtle growth defects were observed. First, root growth was reduced in a repeatedly selfed, homozygous, phospho-deficient line (**Fig. 4A, B**) as compared to P-enabled eS6A^WT-HA^ and eS6B. Second, young eS6A^1′6S>A^ complemented seedlings had pointed first leaves (**Fig. 4C**), a common phenotype in ribosomal protein mutants, including *rps6a* and *rps6b*. The pointed leaf phenotype is not due to the fact that eS6B is missing because *rps6a rps6b* heterozygotes (which contain 50% each of RPS6A and RPS6B) also have the pointed leaf phenotype (Creff et al., 2010; Ren et al., 2012), and eS6A^WT-HA^ plants did not have pointed leaves (**Fig. 4C**). Third, we newly observed that *rps6a* mutants, and to a lesser degree *rps6b* as well, tend to have cotyledons that differ in size. This asymmetry was retained in the eS6A^1′6S>A^ seedlings (**Fig. 4D-E**).

**Figure 4.**
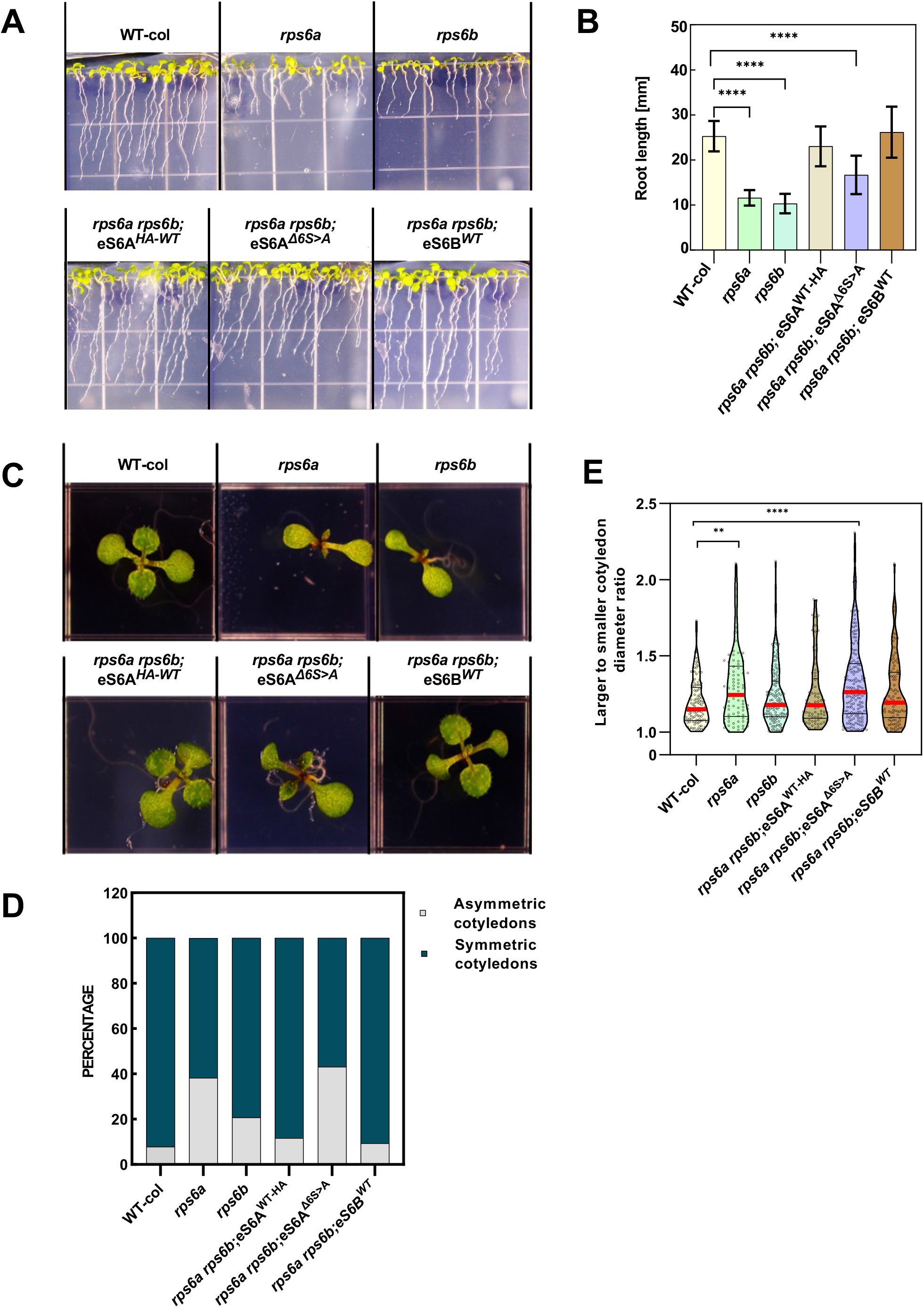
Plants that are deficient for eS6-P grow largely normally with exceptions during seedling establishment. **(A)** Phenotype of 11-day-old seedlings complemented with a phospho-deficient version of eS6A, as compared to controls. **(B)** Root lengths of seedlings from experiment (A). Error bars indicate standard deviations, and **** indicates p<0.0001 by Welch’s t-test. **(C)** Pointed-leaf and asymmetric cotyledon phenotypes in 7-day-old seedlings of P-deficient plants and controls. **(D)** Cumulative histogram and **(E)** violin plot of the asymmetric cotyledon phenotype. Medians are indicated by red bars. ** p=0.0056 and **** p<0.0001 by Mann-Whitney test.

The *rps6a rps6b* double mutants that were complemented with the phospho-deficient eS6A^1′6S>A^ transgene showed essentially normal polysome loading before and after the dark-to-light shift (**Fig. 3C**; quantified in **Supplemental Fig. 7B)**. In these experiments the plant growth or centrifugation conditions differed from the earlier experiments, resulting in numerically lower polysome loading. However, exposure to light increased the polysome loading in both the phospho-enabled eS6A^WT-HA^ and the P-deficient eS6A^1′6S>A^ mutant. Similar results were obtained when the double mutants were complemented with eS6B, or eS6B^1′3S>D^ **(Supplemental Fig. 7A)**. As expected, light-induced phosphorylation of eS6 in polysomes was detected in wild-type plants and in double mutants complemented with eS6A^WT-HA^, but not in plants complemented with eS6A^1′6S>A^ (**Fig. 3D**). Therefore, eS6 protein functions apparently normally in supporting polyribosome loading despite being unphosphorylated (Δ6S>A) or harboring *bona fide* phospho-mimic mutations (Δ3S>D). In addition, in these experiments all the transgenes suppressed the overaccumulation of the 60S subunit **(Supplemental Fig. 7A, B)**.

To characterize the phospho-deficient plants in more detail, we measured the photosynthetic efficiency of photosystem II (Qy_max_ = F_v_/F_m_). In keeping with misexpression of photosynthesis genes in *rps6a* mutant seedlings (*see* RNA-Seq experiment below), PS II efficiency was reduced in *rps6a* and *rps6b* mutants in seedlings (**Fig. 5A-B**) and in rosette-stage plants grown at 22°C or at 12°C (**Fig. 5C-D**). The phospho-deficient plants (*rps6a rps6b*; eS6A^1′6S>A^) were largely rescued for photosynthetic efficiency at all stages.

**Figure 5.**
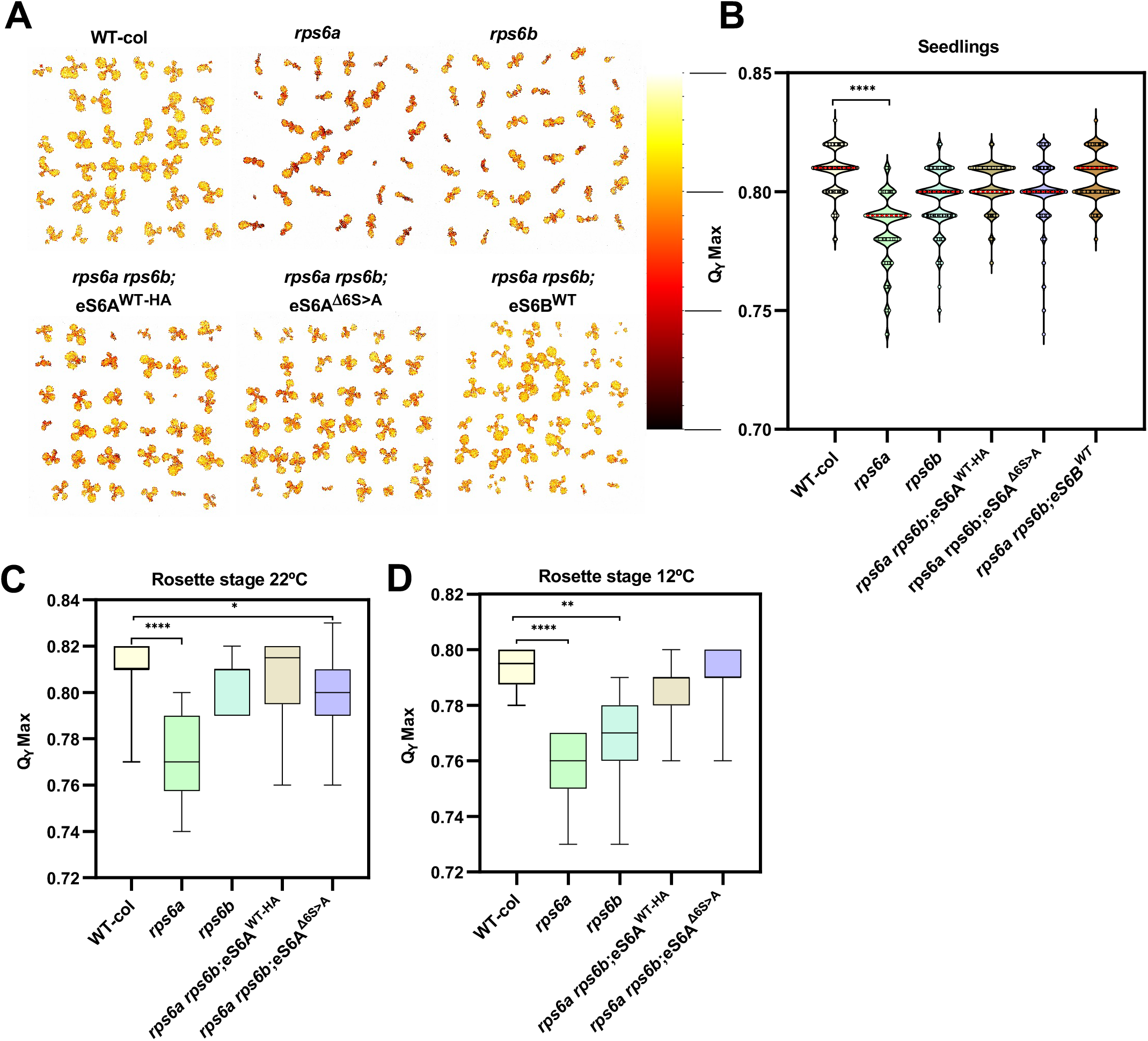
Phenotypes of P-deficient plants in mature rosettes and photosynthetic efficiency. **(A)** Qy_max_ heatmaps in 11-day-old seedlings for the indicated genotypes of P-deficient plants and controls. **(B)** Quantification of data from experiments such as panel (A). **(C, D)** Qy_max_ in 3-week-old rosettes for the indicated genotypes of P-deficient plants and controls at **(C)** 22°C and **(D)** 12°C.

### Gene expression defects in seedlings lacking eS6-phosphorylation

The transcriptome of *rps6a rps6b;* eS6^Δ6S>A^ phospho-deficient seedlings was analyzed in triplicate by RNA-sequencing alongside with eS6A^WT-HA^ complemented plants, *rps6a* and *rps6b* mutants, and wild type. Mapping the reads originating from the *RPS6* genes confirmed that the eS6^Δ6S>A^ plants indeed only contained phosphomutant eS6 (**Supplemental Fig. 8**).

Principal component analysis (PCA) revealed that the three biological replicates clustered closely together, while the five genotypes were distinct. *rps6a* and *rps6b* were clearly distinct from WT in PC1, and also differed from each other in PC2 (**Fig. 6A**). Filtering for differentially expressed genes between WT and the mutants and complementation lines confirmed that eS6^WT-HA^ and eS6^Δ6S>A^ transgenes substantially complemented the *rps6a rps6b* double mutation (**Fig. 6C**). Of 93 mRNAs that were differentially expressed at FDR<0.05 between eS6^WT-HA^ and WT, only 15 differed by >2-fold. The phospho-deficient eS6^Δ6S>A^ transgene complemented the double-*rps6* phenotype slightly less well than the phospho-enabled eS6^WT-HA^ transgene, as is visually evident from the volcano plots, venn diagram, and heat map (**Fig. 6B-D**).

**Figure 6.**
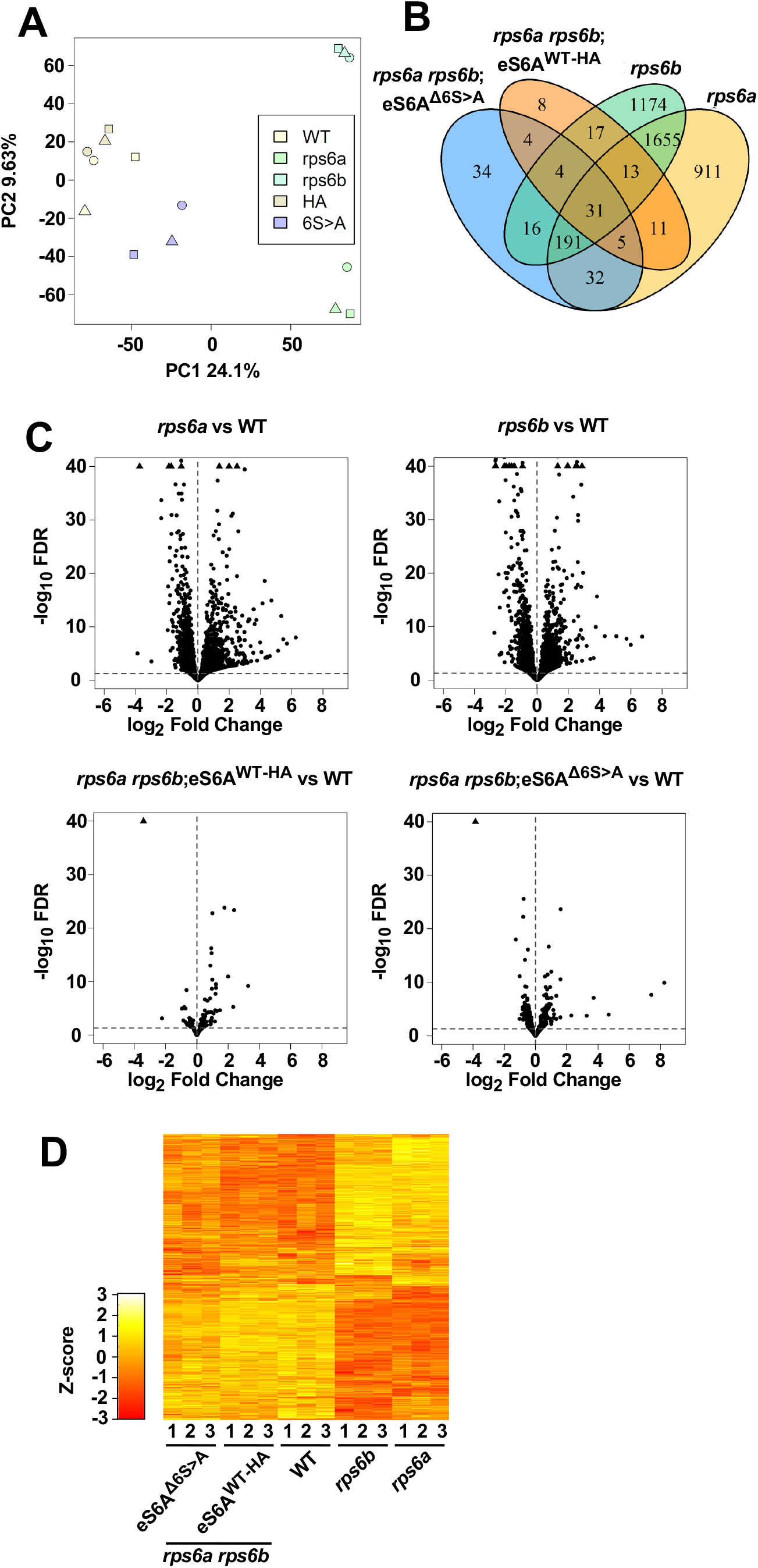
The transcriptome of seedlings with a phospho-deficient eS6A resembles that of wild type, but falls short of full functional complementation in specific ways. **(A)** Principal component analysis of the 15 samples. **(B)** Venn diagram showing the number of differentially expressed genes (FDR< 0.05, versus wild type) in various pairwise comparisons. **(C)** Volcano plots of differential gene expression. The stippled horizontal line marks the false discovery rate (FDR) cutoff of 0.05 that defines differentially expressed genes. LogFC = log_2_ of the fold difference. **(D)** Heatmap of differentially expressed genes. Only genes that passed FDR in at least one of the six relevant pairwise comparisons are included. Genes were filtered with DESeq2, and the display is Z-scored.

The eS6A^1′6S>A^ transgene showed little evidence for gain of function effects, i.e. differential gene expression over wild type that cannot be interpreted as a lack of complementation of the *rps6a* and *rps6b* mutations (34+4=38 genes, i.e. 11% of DEGs between eS6^Δ6S>A^ versus WT, **Fig. 6B**). This number is barely above the number expected for false discovery, and their fold-difference in expression was generally well below two-fold, suggesting they are statistical outliers. If eS6-P played a ‘moonlighting’ role independent of eS6’s role in the ribosome, this role might have revealed itself here in the form of novel aberrations in the gene expression profile, but it basically did not. Instead, the majority (∼90%) of genes that are differentially expressed in eS6A^1′6S>A^ are already known to be misexpressed in the *rps6* mutants. Most likely these residual weak effects in eS6A^1′6S>A^ are loss-of function effects; they may be due to the phosphodeficiency or due to a shortfall in eS6 gene expression (see **Fig. 3B**). This is difficult to distinguish rigorously.

The *rps6a* and *rps6b* mutants have a moderate number of paralog-specific gene expression defects (e.g. see **Fig. 6B, D**). Jointly, the two eS6A-based transgenes were about as successful in complementing *rps6b*-specific defects as *rps6a*-specific defects (**Fig. 6B, D**; all but 4 of the *rps6b*-specific genes and all but 5 of the *rps6a*-specific genes). This again confirms that the eS6A and eS6B proteins are for the most part functionally equivalent.

For the analysis of the eS6^Δ6S>A^ phosphodeficient strain we focused on genes and gene ontology terms that distinguish it from the eS6^WT-HA^ strain. Of 317 genes that passed FDR, only 13 were altered by more than 2-fold (**Supplemental Dataset 1).** Gene ontology analysis with TOPGO revealed that many functional categories (GO terms) were mis-expressed in *rps6a* and *rps6b* versus WT (**Fig. 7**), for example ‘translation’ and ‘photosynthesis’. Gene-by-gene heatmaps demonstrate that the defects in *rps6a* and *rps6b* were qualitatively if not quantitatively the same for most genes (**Supplemental Fig. 9A, B, F**). Remarkably these defects were almost fully complemented by both the eS6A^WT-HA^ and eS6A^1′6S>A^ transgenes. Thus, phosphorylation of eS6A is not required for expression of most translation-related mRNAs. This result is consistent with our polyribosome analyses (**Fig. 6C**). Likewise, both the eS6A^WT-HA^ and eS6A^1′6S>A^ transgenes complemented the *rps6* mutants’ major deficiencies in ‘photosynthesis’, including the major child terms, photosystem I and II, light harvesting, and photoinhibition (**Supplemental Fig. 9B**), consistent with the rescue of photosynthetic efficiency, Qy_max_ (**Fig. 5A-D**).

**Figure 7.**
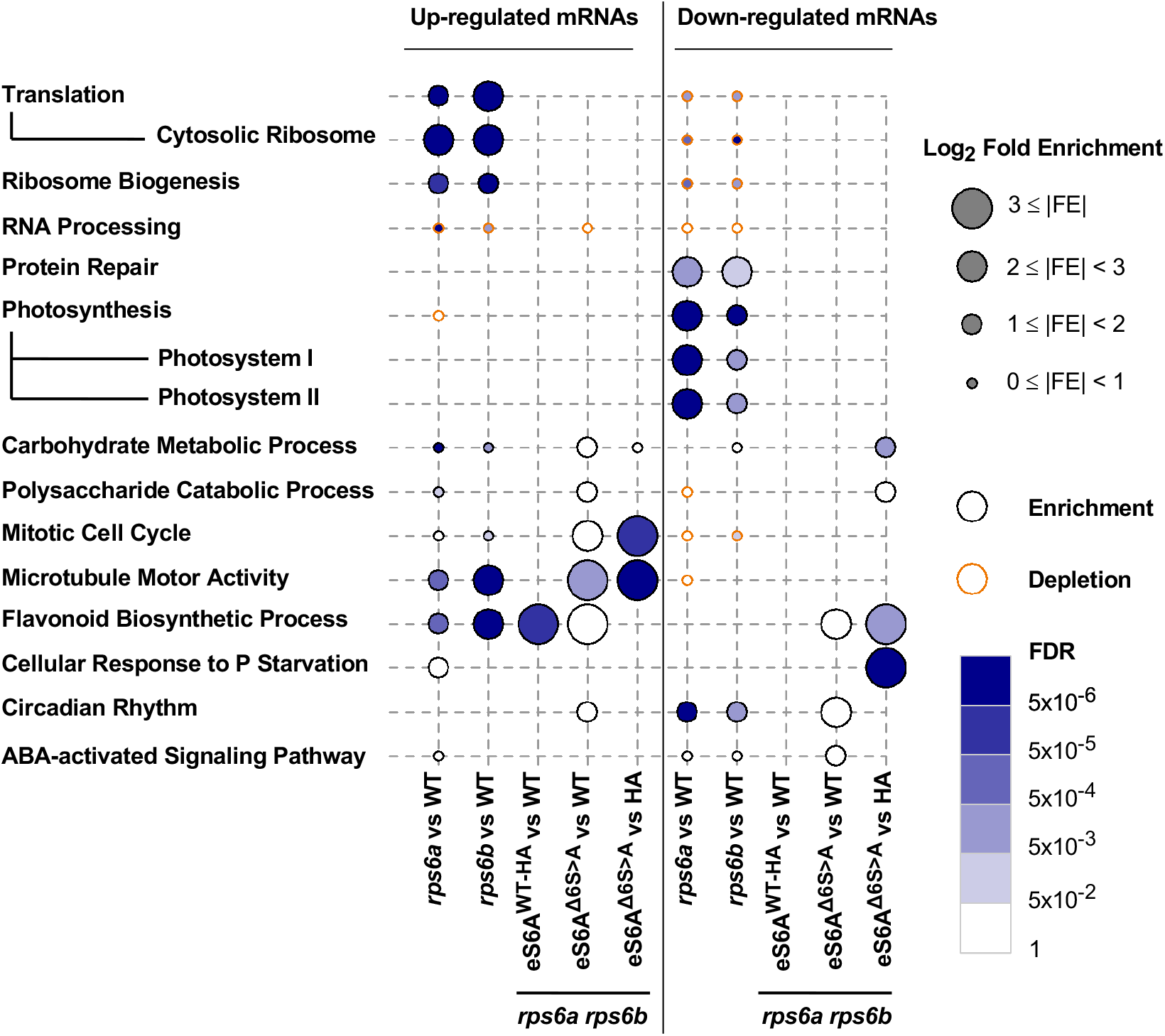
RNA-Seq gene ontology (GO) analysis. The graphic shows functional trends among genes that were differentially expressed between wild type and mutant or transgenic lines. The circle size for each GO term indicates the fold-enrichment of the functional term among the differentially expressed genes over what was expected by chance alone. The fill color indicates the likelihood of false discovery (FDR). The gray perimeter indicates functional enrichment and the orange perimeter indicates depletion.

In contrast, several other GO terms were not fully complemented by eS6A^1′6S>A^. Prominent among these were ‘mitotic cell cycle’, and ‘microtubule motor activity’ (**Fig. 7** and **Supplemental Fig. 9C-D**), which was driven by upregulation of kinesins. The phospho-deficient eS6 appears to be not fully functional as compared to eS6A^WT-HA^ with respect to cell cycle functions.

A function specifically altered between eS6A^1′6S>A^ and eS6A^WT-HA^ was ‘cellular response to phosphate starvation’. These genes were coordinately altered in both *rps6a* and *rps6b*. The eS6A^WT-HA^ slightly <u>over</u>complemented these defects, while eS6A^1′6S>A^ fell slightly short. This caused a notable differential between eS6A^1′6S>A^ and eS6A^WT-HA^ (**Supplemental Fig. 9E**). Of 11 genes significantly underexpressed in the phospho-deficient line, four were involved in galactolipid and sulfolipid synthesis (MGD2, MGDC, SQD1, SQD2), besides *PHO* and *PAH* and *SPX* genes involved in other aspects of phosphate homeostasis. This raises the interesting idea that eS6-P may play a role in phosphate homeostasis.

### Proteome effects in *rps6* single mutants and P-deficient double mutants

To our knowledge no proteome analysis has been published for Arabidopsis ribosomal protein mutants. We performed mass spectrometry proteomics in triplicate on the same five genotypes previously subject to RNA-Seq. Similar to the RNA-Seq data, the PCA separated the five genotypes based on differential protein abundances (**Fig. 8A**).

**Figure 8.**
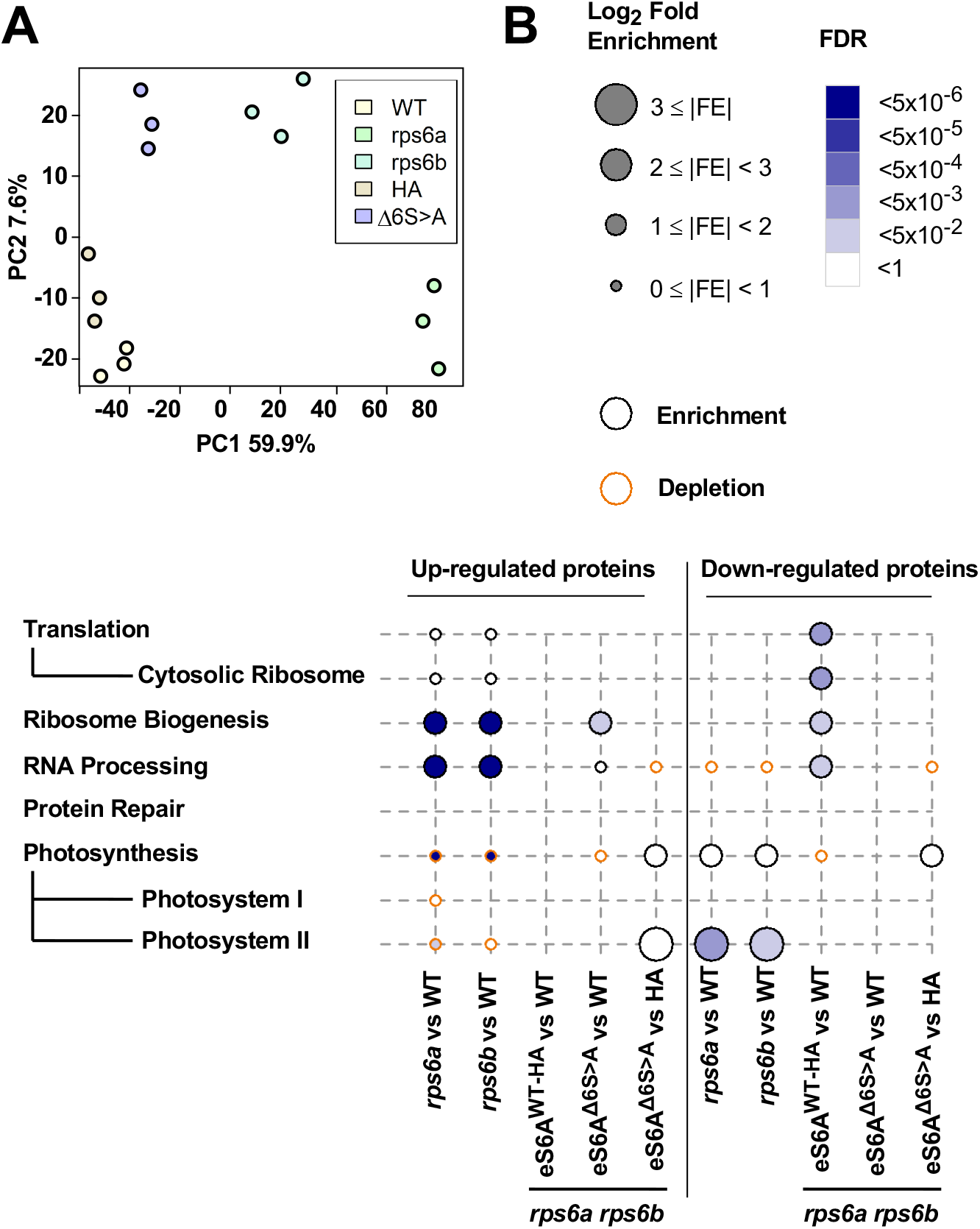
Proteome of *rps6a* and *rps6b* ribosomal protein mutants and double mutants complemented with phospho-deficient and phospho-enabled eS6A. Protein samples are from shoots of ∼12-day-old seedlings. **(A)** Principal component analysis of the triplicate data mirrors the separation of the samples in the RNA-Seq analysis. **(B)** Differentially expressed proteins that passed a significance threshold were classified by gene ontology. Major enriched terms around ribosomes and photosynthesis are displayed with their enrichment or depletion factor and likelihood of false discovery.

The *rps6a* and *rps6b* mutants were deficient in proteins for photosynthesis, especially photosystem II. They overaccumulated ribosome biogenesis proteins, many ribosomal proteins, and other RNA-related proteins. While the deficiency in photosynthesis proteins was rescued effectively by both phospho-enabled and phospho-deficient eS6A (**Fig. 8B and Supplemental Figure 10C**), interestingly, the excess in ribosomal biogenesis proteins was rescued less well by phospho-deficient eS6A^1′6S>A^ than phospho-enabled eS6A^WT-HA^. This defect for ribosome biogenesis was more pronounced than that for ribosomal proteins *per se* and for other RNA-related functions such as RNA degradation and translation initiation (**Fig. 8B and Supplemental Figure 10A, B and D**). Complementation of photosynthesis and translation defects were both consistent with the RNA-Seq data. These results may suggest that the ability to phosphorylate eS6A represses the mRNA expression or accumulation of proteins for ribosome biogenesis. A role for eS6-P in ribosome biogenesis had previously been suggested in the mouse (Chauvin et al., 2014), although it played out at the transcriptional level rather than translation.

## DISCUSSION

This analysis was motivated by the question as to the functional significance of the phosphorylation in the carboxy-terminal tail of the ribosomal protein eS6.

### New phenotypes of ribosomal protein mutants

We adopted a new T-DNA insertion allele for the *RPS6A* paralog, *rps6a-2*, and found that it is a null allele for both mRNA and protein that recapitulates all the phenotypes examined of the earlier allele, *rps6a-1* (Creff et al., 2010). Our analysis discovered several new phenotypes of *rps6* mutants. The two cotyledons of *rps6* mutants often differ in size. The pale green color of the seedlings and rosette-stage plants is accompanied by defects in gene expression of many photosynthetic genes and by reduced quantum efficiency of photosystem II, an effect exacerbated at cooler temperature. Our data demonstrate the detrimental effects of cytosolic ribosomal protein mutations on the chloroplast at the level of the transcriptome, proteome and photosynthetic efficiency. The *rps6a* and *rps6b* mutants have an excess of free 60S subunits, which was observed before in yeast and Arabidopsis (Pachler et al., 2004; Creff et al., 2010), but is documented here with statistical support. We also provide one of the first transcriptome datasets from mutants in plant cytosolic ribosomal proteins, and to our knowledge the first that compares mutations in multiple paralogs. The *rps6a* and *rps6b* mutant transcriptomes are largely similar but differ in the details. In *rps6a* the photosynthesis functions are more strongly affected than in *rps6b*, while in *rps6b* the cytosolic translation functions are more strongly upregulated than in *rps6a*. The plastid ribosomes don’t feature prominently. Because transgenes of eS6A that use the *RPS6A* promoter complemented both functions well, we suspect that the difference in phenotype is not due to differences in the amino acid sequences of eS6A and eS6B. Instead, this difference could arise if eS6B had a shifted spatial expression pattern, biased towards growing cells, which produce more ribosomes, rather than cells fated to become mesophyll, which produce the photosynthetic apparatus.

We also present the first proteomics data for plant ribosomal protein mutants. Defects include a drop in photosynthesis functions and an upregulation of ribosomal proteins and ribosome biogenesis proteins in both *rps6a* and *rps6b*, consistent with the transcriptome data. The *rps6a* mutant has strongly elevated protein levels of a subset of ribosomal proteins and even more strongly for Ribi proteins. This same effect is also seen in *rps6b*, mostly for the same ribosomal proteins. In each mutant the other paralog is upregulated, together with many other ribosomal proteins. Together, these results suggest the mutants’ growth is rate limited by production of *rps6* mRNA. In the *rps6b* mutant it takes arguably twice as long to accumulate enough *RPS6* mRNA per cell than in wild type. Hence the mutants grow more slowly. Meanwhile, they accumulate excess mRNAs for other ribosomal proteins, whose gene dosage is normal, and which overaccumulate as a result. Ribosome biogenesis proteins are expressed at a higher level as well, perhaps driven by the excess of yet to be assembled ribosomal proteins.

### eS6A and eS6B paralogs are functionally equivalent

We looked for evidence that the A and B paralogs of eS6 have different functions. In short, our data suggest that the two paralogs are functionally equivalent. First, the eS6B paralog was able to complement the *rps6a* mutant phenotype as well as the eS6A paralog, with respect to root growth and general appearance, as observed earlier (Creff et al., 2010; Ren et al., 2012). Second, the double mutant *rps6a rps6b* was complemented effectively by eS6A^WT-HA^, a hemagglutinin-tagged wild-type of eS6A, as well as a wild-type version of eS6B. Even though both wild-type alleles were absent, both eS6A and eS6B were able to complement essentially completely and equally. The phenotypic rescue was evident at the biochemical level (polysome profiles and loading), photosynthetic efficiency, root growth, and other developmental phenotypes. We note here that the single mutant *rps6b* was complemented incompletely by either eS6A or eS6B, a result that we hypothesize may be due to residual mRNAs from the *RPS6B* gene interfering with transgene expression. In keeping with this explanation, in the *rps6a rps6b* double mutants, complementation with wild type *RPS6* genes was complete, and coincidentally, the residual transcripts from the mutated *rps6b* gene were suppressed.

### Phospho-deficient eS6 performs many functions of the wild-type eS6

Light, sucrose and auxin can boost translation. These signals also boost TOR activity, which then causes phosphorylation of eS6 in its C-terminal tail. This has led to the hypothesis that eS6-P is causally involved in mediating these effects on translation. However, evidence for this idea has been mostly correlative (Boex-Fontvieille et al., 2013; Dobrenel et al., 2016; Chen et al., 2018). For example, eS6-P is accompanied by increased expression of plastid ribosomal proteins (Dobrenel et al., 2016) and by increased polysome loading in the cytosol (Liu et al., 2012; Chen et al., 2018); however, our data indicate that eS6-P-deficient plants express chloroplast ribosomal proteins about normally, and can boost polysome loading in the morning as effectively as wild type. Recently, Mancera-Martinez et al. addressed this question by manipulating the phosphorylation potential of eS6 by expressing ΔS>A and ΔS>D versions of eS6B from a cauliflower mosaic virus 35S promoter in *rps6a* single-mutant plants. Similar to our data, the two versions did not differ in global polysome loading. However, the two versions differed in the translation of two reinitiation-dependent reporter mRNAs, suggesting that eS6 phosphorylation may regulate the translation of specific mRNAs (Mancera-Martinez et al., 2021).

Here, we sought out evidence to reveal the role of eS6 phosphorylation by comparing the biochemical, molecular, cellular, and organ-level phenotypes of *rps6* double mutants complemented with either a phosphorylation-enabled allele (WT-HA) or a phosphorylation-deficient allele (Δ6S>A). This is the first study to examine P-deficient alleles in a genetic background mutated for both *RPS6* paralogs. It is also the first to compare the P-deficient eS6 against the wild type, rather than P-deficient against P-mimic (Mancera-Martinez et al., 2021). We chose the eS6A paralog, rather than both eS6A and eS6B for simplicity. This was a reasonable choice because eS6A and eS6B were confirmed to be functionally equivalent. We constructed the phospho-deficient allele as close to the native version as possible. Aside from keeping its native promoter, untranslated regions, and exon-intron structure, we left it untagged in order to not introduce confounding features. We did tag the corresponding wild-type eS6A with HA to be able to distinguish it from the native version, but our data suggest that this did not compromise its function in a major way. We created a series of phospho-deficient versions of eS6A and eS6B anticipating that this would allow us to define the minimal changes necessary for a robust phenotype. In the end, a version with six codons altered to alanine revealed only rather subtle phenotypes, and therefore we did not pursue the versions with fewer changes in detail. However, data from other S>A and S>D alleles are included in this article because their results reinforce those for the main Δ6S>A allele.

Overall, the phospho-deficient eS6A proved to be largely functional, given that it reverted the lethal phenotypes of the *rps6a rps6b* double mutant back to normal. In detail, polysome loading was normal, including the boost in polysome loading during the daily dark-to-light transition. And while *rps6* single mutants had an excess of the 60S subunit, the phospho-deficient double mutants reverted back to normal. The phospho-deficient eS6A also partitioned to cellular compartments, nucleus, nucleolus, and cytosol, similar to wild type. And it rescued most gene expression defects commonly seen in *rps6a* and *rps6b* mutants, as well as misregulation of protein levels in translation and photosynthesis. Accordingly, photosynthetic efficiency was mostly back to wild-type levels.

Here we want to emphasize that our conclusion is not solely based on the *rps6a rps6b* double mutants. Even in *rps6a* single mutants, surprisingly, the presence of a phospho-deficient eS6A suppressed the phosphorylation of the eS6B paralog, yet rescued the polysome profiles of *rps6a* and allowed for normal growth. Although we can only speculate about the mechanism for this coordinated loss of phosphorylation, this result bolsters the conclusion that phospho-deficient eS6 is largely functional, and ribosomes lacking eS6 phosphorylation are functional as well.

### Emerging roles of eS6 phosphorylation

Detailed analysis of the phospho-deficient plants did, however, reveal several notable abnormalities and deficiencies. The YFP-tagged eS6A^1′7S>A^ tended to aggregate a bit more readily in the nucleus than did the wild-type version. In the gene expression profile, certain defects in the mitotic cell cycle functions and responses to phosphate starvation, which are characteristic of *rps6* mutants, failed to get fully rescued by the phospho-deficient Δ6S>A allele, although they were rescued by the phospho-enabled WT-HA allele. A transient pointed-leaf phenotype suggested that leaf expansion was slightly delayed, as was the rate of root elongation, a sensitive indicator of cell division. At the proteome level, certain misregulations of ribosome biogenesis proteins also failed to get fully rescued. In this context it is notable that eS6 functions as part of the small subunit processome in ribosome biogenesis (Bernstein et al., 2004). The increase in ribosomal proteins and the excess free 60S subunit seen in our *rps6* mutants mirror similar increases in yeast and mammalian ribosomal protein mutants (Pachler et al., 2004; Robledo et al., 2008).

Together, these results are our best indicator as to the cellular function of eS6-phosphorylation in Arabidopsis. The phospho-deficient plants had morphological defects as seedlings, cotyledon asymmetry and pointed leaves. Although these phenotypes were more transient than in the original *rps6* mutants, meaning they had largely disappeared at the time when RNA was harvested for transcriptome profiling, it is plausible that it was the misregulation of cell cycle related mRNAs and ribosome biogenesis proteins that led to the growth inhibition in cotyledons and first leaves.

It was plausible that the eS6 phospho-deficient plants would reveal a phenotype not seen in *rps6* mutants, a gain-of-function defect. Notably, few if any of the defects observed in the phospho-deficient plants look like a clear gain-of-function defect. Instead, the phenotypes in the *rps6a rps6b* eS6A^1′6S>A^ line are incomplete rescues of defects germane to *rps6a* mutants. Are there not at least a few individual genes that show the gain-of-function pattern, i.e. abnormal expression in Δ6S>A, but normal expression in all other genotypes? Yes, but very few genes (34 by one estimate) match this pattern. These include ADC2 and RD19, CR88 and CYP707A2, CHIL and RHM1, and BBD1. These genes may be the rare sentinels of eS6-phosphorylation, but this inference is not strong, because these genes may also be false-negatives for differential expression in the *rps6a* and *rps6b* mutants, and they are not legitimized by belonging to a functionally defined gene ontology group.

This study has a number of limitations. Although the phospho-deficient version of eS6A was expressed at a level higher than the native eS6A, it did not quite reach the two-fold increase one would have preferred. Our proteome data in the *rps6a rps6b* background pegged eS6A^1′6S>A^ at 1.6-fold the level of eS6A in wild type. Because *RPS6A* and *RPS6B* are expressed equally, the total level of transgenic eS6 protein was 20% below wild type. We consider this acceptable because *rps6a* heterozygotes, whose gene dosage is 25% below wild type, do not have major phenotypes, i.e. *rps6a* and *rps6b* are recessive mutations, as is also true for other *rps* mutations (Van Lijsebettens et al., 1994). Our work also did not check for phenotypes under stress conditions, with photosynthesis at 12°C as one exception. It is possible that the P-deficient plants might reveal additional growth defects under conditions that alter eS6-P, such as hypoxia, heat, or light and darkness.

Taken together, our data together with those of Mancera-Martinez and coworkers (Mancera-Martinez et al., 2021) extend what we know about the functional consequences of eS6-P to a photosynthetic organism, Arabidopsis. In each eukaryote that has been studied, yeast, the mouse, and now Arabidopsis, global translation is largely unaltered (Ruvinsky et al., 2005; Yerlikaya et al., 2016), although Wittenberg and coworkers reported a lower rate of translation and increased translation fidelity in eS6-P-enabled cells (Wittenberg et al., 2016). Evidence for effects on mRNA-specific translation exists but is sparse (Puighermanal et al., 2017; Bohlen et al., 2021; Mancera-Martinez et al., 2021). This is also true for effects at the transcriptome and proteome level ((Yerlikaya et al., 2016) and this work). Translatome defects were not examined in our study. These minimal phenotypes stand in contrast to the diverse consequences of the environmental stimuli that regulate eS6-P and the diverse phenotypic effects from inhibiting the TOR-S6 kinase pathway genetically or pharmacologically (Ren et al., 2012; Ahmad et al., 2019; Margalha et al., 2019; Scarpin et al., 2022; Ingargiola et al., 2023). Yerlikaya and coworkers (Yerlikaya et al., 2016) concluded conservatively that none of their phenotypes in eS6 P-deficient yeast could be firmly ascribed to the P-deficiency but were probably due to reduced eS6 protein levels. For this work in Arabidopsis we like to propose a similar caveat. Essentially all the phenotypes seen in eS6 phospho-deficient plants are also seen in *rps6* mutants. They are hypomorph (loss-of-function) phenotypes rather than neomorph (gain-of function) phenotypes. It is therefore difficult to rule out that these phenotypes are due to the reduced level of the eS6A^1′6S>A^ protein or subtle insufficiencies in its pattern of expression, rather than its lack of phosphorylation.

## MATERIALS AND METHODS

### Arabidopsis strains and genotyping

For *RPS6A* (At4g31700), the *rps6a-2* allele is line GK_468C04 from the GABI-KAT T-DNA collection with an insertion in intron 4 (Kleinboelting et al., 2012). For *RPS6B* (At5g10360) the *rps6a-1* allele is SALK_012147C from the SALK collection with an insertion in exon 4 (Creff et al., 2010).

*rps6a-2* and *rps6b-1* seedlings were genotyped via PCR to check for the presence of the WT RPS6 gene and the T-DNA insertion. LP and RP primers that span the T-DNA insertion site were used to amplify the WT fragment which is ∼1kb in size. LP and T-DNA LB (GabiKat) primers were used to amplify the T-DNA region which is ∼0.7 kb. Similarly, for *rps6b-1* seedlings, LP and RP primers were used to amplify the WT fragment and RP and TDNA LBb 1.3 (SALK) primers were used to amplify the T-DNA fragment. Annealing temperatures of 56°C and 54.3°֯C were used for *rps6a-2* and *rps6b-1* reactions respectively.

To analyze segregation of the transgenes seedlings were grown on 1⁄2 strength MS medium supplemented with 1% sucrose with either 50 μg/ml kanamycin, 7 μg/ml sulfadiazine or Basta. 50 seeds were plated on either kanamycin or sulfadiazine to score the 3:1 segregation of *rps6b-1* or *rps6a-2*, respectively, and 70 seeds were plated on Basta plates to score for a 3:1 or 15:1 or 63:1 segregation of the transgene. Resistance versus sensitivity was scored 7 days after plating, and a Chi-square test was performed to confirm the segregation ratio. The line was then planted in soil and the progeny from the plants was again scored the same way on the way to identify families homozygous for *rps6a-2*, *rps6b-1* and a transgene.

### RNA and protein methods

For reverse-transcription PCR, WT, *rps6a-2* and *rps6b-1* seedlings were grown on 1⁄2 strength MS salt medium for 12 days and the tissue was harvested for RNA extraction using the Zymo Research ZR Plant RNA Mini Prep Kit. cDNA was synthesized from the total RNA, and the *RPS6A* and *RPS6B* transcripts were amplified via PCR using PrimeStar Max polymerase (Takara). The PCR products were separated on a 1% agarose gel. SDS-PAGE and immunoblotting with phosphospecific antibodies were performed as described (Enganti et al., 2018).

### Polysome Profiling

The aerial portion of 12 days-after-germination seedlings grown under long day conditions was collected by flash freezing at ZT23.5 (30 minutes before lights on) and ZT2.5 (two and a half hours after lights on). Plant tissue was ground in liquid nitrogen and extracted in polysome isolation buffer (200 mM Tris-HCl pH 8.4, 25 mM MgCl_2_, 50 mM KCl, 1% deoxycholic acid and 2% polyoxyethylene 10 tridecyl ether). Sucrose gradients were prepared by layering 1.7 ml, 3.3 ml, 3.3 ml, and 1.7 ml each of 50% sucrose, 38.4% sucrose, 26.6% sucrose, and 15% sucrose, respectively. After addition of each gradient layer, the centrifuge tube was frozen at -80°C for 1 h. On the day before use, the gradients were thawed overnight without shaking at 4°C. Plant extracts (1 ml) were loaded on top of a 10 ml 15–50% sucrose gradient and centrifuged at 35,000 rpm for 3.5 h without brake (Beckmann Coulter SW 41Ti). After recording the RNA absorbance profile at 254 nm, the gradient was fractionated into 12 equal fractions. Samples from the fractions were then separated on SDS–PAGE gels followed by immunoblotting to determine eS6 protein levels and phosphorylation levels in the samples. Equal volumes of sample from each fraction were loaded onto the gels.

Area under the curve for the polysome profile traces were calculated as described (Enganti et al., 2018; Lokdarshi et al., 2020b). In brief, gradient traces were manually split into 40S, 60S, 80S, small polysomal (2-4 ribosomes) and large polysomal sections (5+ ribosomes). Blank gradient traces were subtracted from the sample traces and the area under the curve was calculated for each section. If applicable, areas were combined into non-polysomal (sum of areas from 40S, 60S and 80S) and polysomal (sum of areas from small and large polysomal) sections. Abundances of different ribosomal complexes were compared between genotypes or time points by using Welch’s t test.

### Transient gene expression in *Nicotiana benthamiana*

Coding sequences of eS6 were cloned as N-terminal fusions to enhanced yellow fluorescent protein and expressed from the CaMV 35S promoter. Overnight cultures of *Agrobacterium tumefaciens* harboring the T-DNA plasmid of interest were grown with appropriate antibiotics. The cultures were pelleted and resuspended in infiltration buffer (10mM MES buffer, 10mM MgCl_2_, pH 5.4) to an optical density of 1.0 at 600nm. Acetosyringone was added to a final concentration of 200μM to the culture and incubated with agitation for 2h. Young leaves on three-week-old *N. benthamiana* plants were infiltrated with the cultures using a 1ml syringe. The plants were kept in the dark overnight and shifted to normal growth conditions for 36h following which the leaves were imaged to detect fluorescence by confocal laser scanning microscopy.

### Root phenotyping

To measure root lengths, seedlings were grown vertically on square petri plates with a grid and photographed on day 7 after germination. Images were imported into ImageJ and root lengths were measured by tracing each primary root using the segmented line tool.

### Photosynthetic Efficiency Measurement

The maximum quantum yield of photosystem II (PS II) (Qymax = F_v_/F_m_) was measured on a FluorCam 800MF (Photon Systems Instruments, Drásov, Czechia) as per the manufacturer’s instructions and modifications (Murchie and Lawson, 2013). Briefly, plants were dark adapted for 1 min (F_0_) prior to applying a saturating pulse of 1800 μEin m^−2^ s^−1^ for 0.8 s (F_m_). Variable fluorescence (F_v_) was calculated as the difference between F_0_ and F_m_ to get the maximum quantum yield (F_v_/F_m_) (Lokdarshi et al., 2020a).

### Molecular cloning and site directed mutagenesis

RPS6A and RPS6B were amplified from WT Col-0 genomic DNA using primers with added restriction sites; *Eco*RI and *Xba*I for *RPS6A* and *Sbf*I and *Pvu*I for *RPS6B*. The amplified product was approximately 3kb which included the full length transcribed region, 1.5kb upstream of the 5’ UTR and 200 bases downstream of the 3’ UTR. Both *RPS6A* and *RPS6B* fragments were digested and then ligated to the T-DNA vector pFGC19 (Kim et al., 2007) that was previously digested with the appropriate enzymes. Site-directed mutagenesis was done by PCR with mutagenic oligonucleotide primers in pFGC19. Serine and threonine codons were mutated sequentially to either alanine (GCT) or aspartate (GAT).

Initially, two fragments were generated by PCR with a single codon substitution in the forward and the reverse strand. The primers that were used generated PCR 1 and PCR 2 fragments of 800 bp and 500 bp for *RPS6A* and 1200 bp and 500 bp for *RPS6B* respectively. The products from PCR 1 and 2 were subsequently mixed to serve as template for PCR 3 to generate a longer fragment with the desired mutation that could then be ligated to pFGC19 harboring the respective RPS6 gene. The PCR 3 products were 1.3 kb for RPS6A and 1.7 kb for RPS6B. The PCR 3 products were purified using a DNA cleanup kit and then digested with either *Bst*BI or *Xba*I for RPS6A or *Xho*I and *Pvu*I for *RPS6B*. *Bst*BI and *Xho*I are internal sites within the coding region of RPS6A (between intron 3 and 4) and RPS6B (exon 2) respectively. *RPS6A* fragments were digested for 30 minutes at 65 C (*Bst*BI) followed by 30 minutes at 37 C (*Xba*I) whereas *RPS6B* fragments were digested at 37 C for 1 hour. The digested products were then run on a gel and purified using a gel extraction kit. The digested fragments were then ligated to pFGC19 digested with the appropriate enzymes in a 3:1 and 1:1 molar ratio of the insert to the vector along with a control that had the cut vector and no insert. The ligation was carried out for 2 hours at 16 C. The ligation products were then transformed into competent Top10 cells via heat shock. The culture was plated on LB plates containing kanamycin and incubated overnight. Colonies were then grown for plasmid extraction and the mutations were confirmed by DNA sequencing. Wild type and mutant T-DNAs were transformed into *rps6a-2 rps6b-1* double heterozygote plants that had been generated by genetic crossing and transgenic seedlings selected for Basta resistance.

### RNA-Seq library construction and sequencing

RNA-seq was performed on WT, *rps6a-2, rps6b-1, rps6a-2 rps6b-1;* eS6A^WT-HA^, and *rps6a rps6b;* eS6A^1′6S>A^ genotypes grown under long day conditions. In the morning 12 days after germination, the aerial portion of the seedlings were harvested by flash freezing in liquid nitrogen. Total RNA was extracted using a commercial kit. RNA quality was measured using a Bioanalyzer (Agilent). Paired end cDNA libraries were constructed using the Illumina Stranded Total RNA Prep with Ribo-Zero Plus. The libraries were sequenced on a NextSeq in paired end mode and with 75 base pair long reads at the Oklahoma Medical Research Foundation (Oklahoma City, USA). Raw read quality was assessed with FastQC v0.11.5. Raw reads were aligned to the TAIR10.1 genome and Araport11 annotation using STAR-2.7.7a (Dobin et al., 2013), with default parameters except for the following: -alignIntronMax 1000. Mapping quality was assessed with RSeQC v4.0.0 (Wang et al., 2012). Reads were counted using subread featureCounts v2.0.1 (Liao et al., 2014) in paired end mode.

### RNA-Seq differential gene expression

Differential gene expression was performed in R (v3.6.3). Genes that were not expressed in all three replicates of at least one sample were removed. Samples were inspected for batch effect by principal component analysis, and no batch effect was found. The filtered reads were then normalized, and pairwise comparisons between genotypes was performed using DESeq2 v1 .26.0 (Love et al., 2014). The resulting p-values were corrected for multiple comparisons using false discovery rate (FDR) with Benjamini-Hochberg parameters and the resulting log2 fold changes were shrunk using ashr2.2 (Stephens, 2017).

Gene ontology analysis was performed on each pairwise comparison using a custom wrapper around the topGO package version 2.38.1; (Alexa and Rahnenführer, 2016). Only genes measured as expressed were used as the gene universe. topGO was run with node size 1, and FDR P value adjustment using a custom script and the classic Fisher, parent-child, and weight01 algorithms. Packages were obtained from CRAN or Bioconductor version 3.7; (Huber et al., 2015).

### Proteomics

Protein digestion was performed as previously described (Enganti et al., 2018). In brief, samples were suspended in a detergent lysis buffer (2% sodium dodecyl sulfate and 10 mM dithiothreitol in 100 mM ammonium bicarbonate) supplemented with Halt Phosphatase Inhibitor Cocktail (Thermo Fischer Scientific) for crude protein extraction. Cell debris was removed, and proteins were alkylated with iodoacetamide (30 mM) and incubated in the dark at room temperature for 15 min. Proteins were precipitated via methanol/chloroform/water precipitation and protein pellets were washed twice with methanol. Dried proteins pellets were resuspended in 1 mL of 8 M urea and incubated at room temperature for 1 hr. Samples were digested via addition of two aliquots of sequencing-grade trypsin (Promega, 1:50 (w:w)) at two different sample dilutions, 4 M urea (overnight) and subsequent 2 M urea (5 h). Following digestion, samples were adjusted to 1% formic acid and desalted using solid-phase C18 extraction cartridges (Sep-Pak Plus Short, Waters), and lyophilized. All samples were analyzed on a Q Exactive Plus mass spectrometer (Thermo Scientific) coupled with an Easy-nLC 1200 (Thermo Fischer Scientific). For each sample, a single 1µg injection of peptides were separated on an in-house-pulled nanospray emitter of 75μm inner diameter containing 25 cm of Kinetix C18 resin (1.7 µm, 100 Å, Phenomenex) across a linear organic gradient of 0–22% (80% acetonitrile, 0.1% formic acid) over 210 min at 200 nL/min. Mass spectra data were acquired with the Thermo Xcalibur software using the top 10 data-dependent acquisition. All MS/MS spectra collected were processed in Proteome Discoverer version 2.2 with MSAmanda (Dorfer et al., 2014) and Percolator (Kall et al., 2007). The spectra were searched against the UniProt reference proteome (Proteome ID UP000006548) to which common laboratory contaminants were appended. The following parameters were used by MSAmanda to derive fully tryptic peptides: MS1 tolerance = 5 ppm; MS2 tolerance = 0.02 Da; missed cleavages = 2; Carbamidomethyl (C, + 57.021 Da) as static modification; and oxidation (M, + 15.995 Da) and carbamylation (N-terminus, + 43.006 Da) as dynamic modifications. The Percolator FDR threshold was set to 1% at the peptide-spectrum match and peptide levels. Pairwise t-tests were performed between protein abundances using Proteome Discoverer. Changes in protein abundances were considered significant with a p-value < 0.05 and a Log_2_ difference > 1.

### Accession numbers

The Arabidopsis locus identifier for *RPS6A* is At4g31700, and that for *RPS6B* is At5g10360. RNA-Seq data are deposited in NCBI-GEO under accession number GSE222967. All proteomics spectral data in this study were deposited at the ProteomeXchange Consortium via the MASSIVE repository (https://massive.ucsd.edu/).

## SUPPLEMENTAL MATERIALS

**Supplemental Figure S1.**
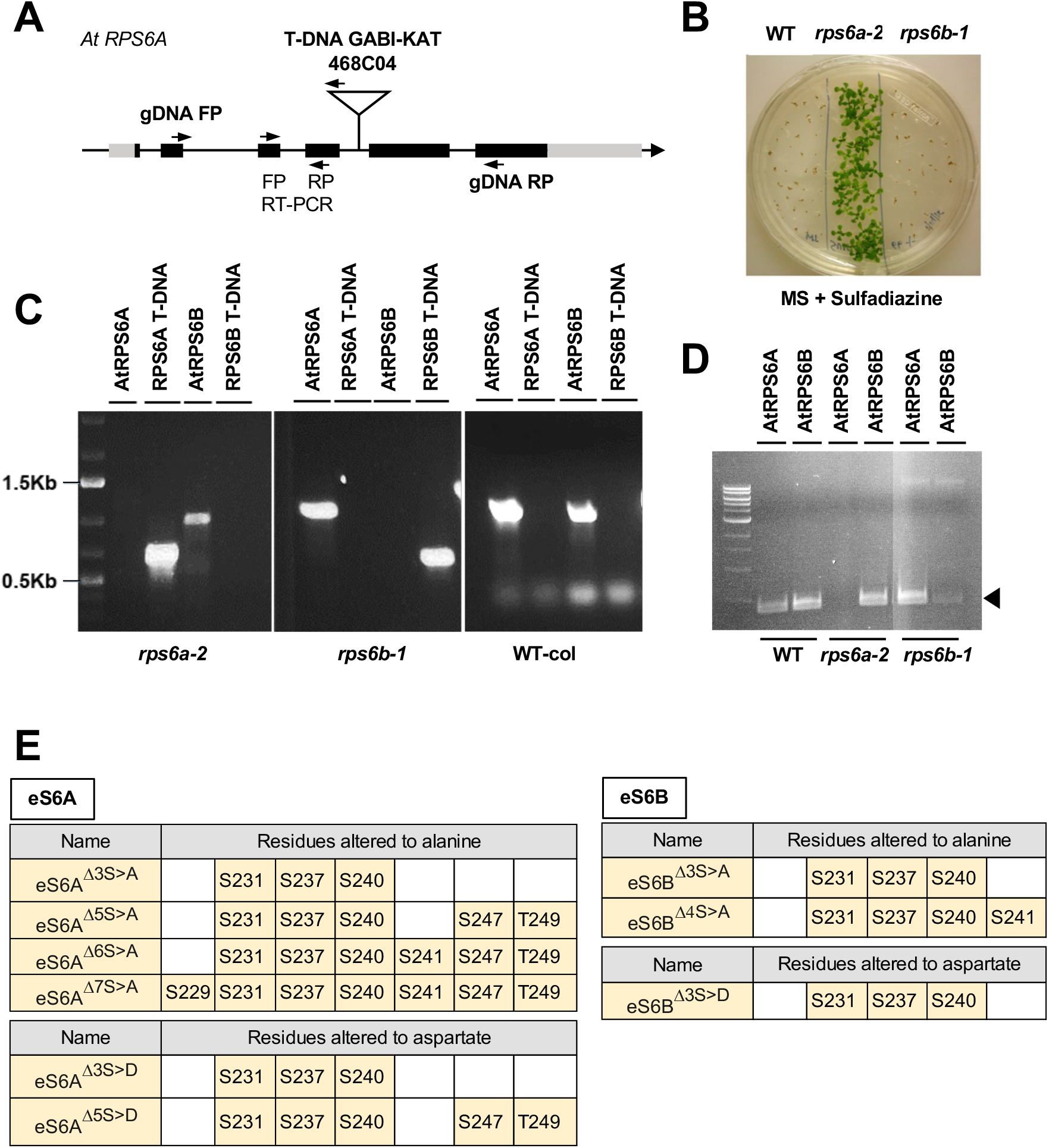
A new null-allele of *RPS6A*, *rps6a-2* is marked by sulfadiazine resistance. **(A)** Gene model showing the T-DNA insertion site in *rps6a-2* (GabiKat line 468C04). Black boxes and gray boxes represent coding and non-coding exons and lines represent introns. Annealing sites of genotyping primers and RT-PCR primers are indicated. **(B)** The *rps6a-2* allele confers resistance to sulfadiazine (7.5 µg/mL). **(C)** Gel image with PCR-genotyping results for *rps6a-2*, *rps6b-1*, and wild-type Columbia seedlings. The PCR scores the presence of the wild-type allele and the T-DNA allele for both RPS6A and RPS6B. **(D)** The *rps6a-2* allele lacks detectable transcript for eS6A. RT-PCR was performed with RNA from WT, *rps6a-2* and *rps6b-1* mutants to determine transcript levels of both *RPS6A* and *RPS6B*. Residual transcripts for *RPS6B* in the *rps6b-1* allele are as expected (Creff et al., 2010). **(E)** Phosphorylation-deficient mutations introduced into eS6A and eS6B.

**Supplemental Figure S2.**
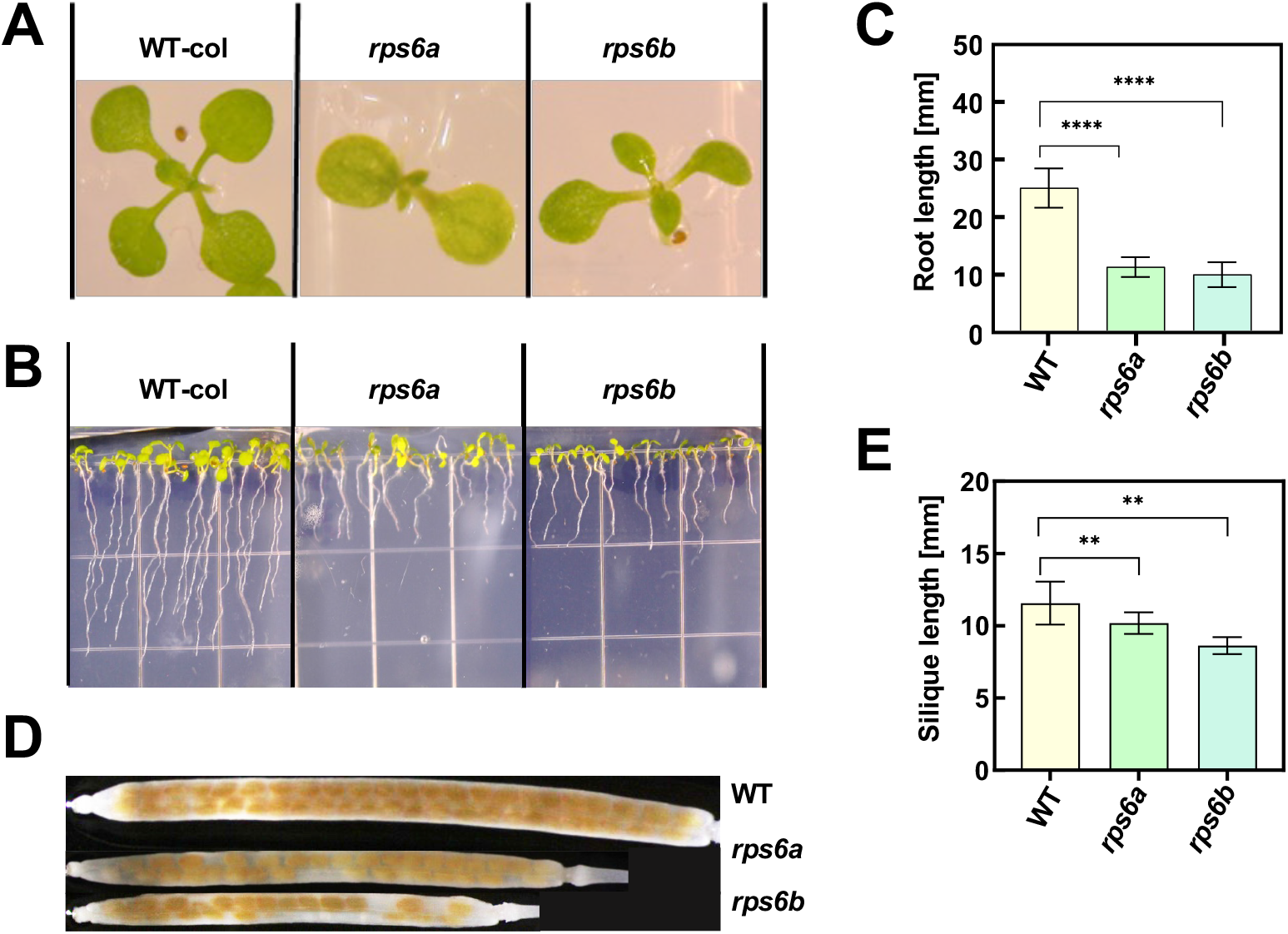
The *rps6a-2* allele recapitulates the growth phenotypes of an earlier allele, *rps6a-1 (Creff et al., 2010)* **(A)** The 10-day old *rps6a-2 and rps6b-1* mutants have pointed leaves compared to WT. **(B)** Root growth of *rps6a-2* and *rps6b-1* mutants is dramatically reduced compared to WT. Images show 7-days-old seedlings. **(C)** Quantification of root lengths. **** indicates p<0.001 by t-test comparing each mutant to WT. Error bars indicate standard error of the mean (S.E.M.). **(D)** Mature siliques of all three genotypes. **(E)** *rps6a-2* produces fewer seeds per silique than wild-type, as does *rps6b-1*.

**Supplemental Figure S3.**
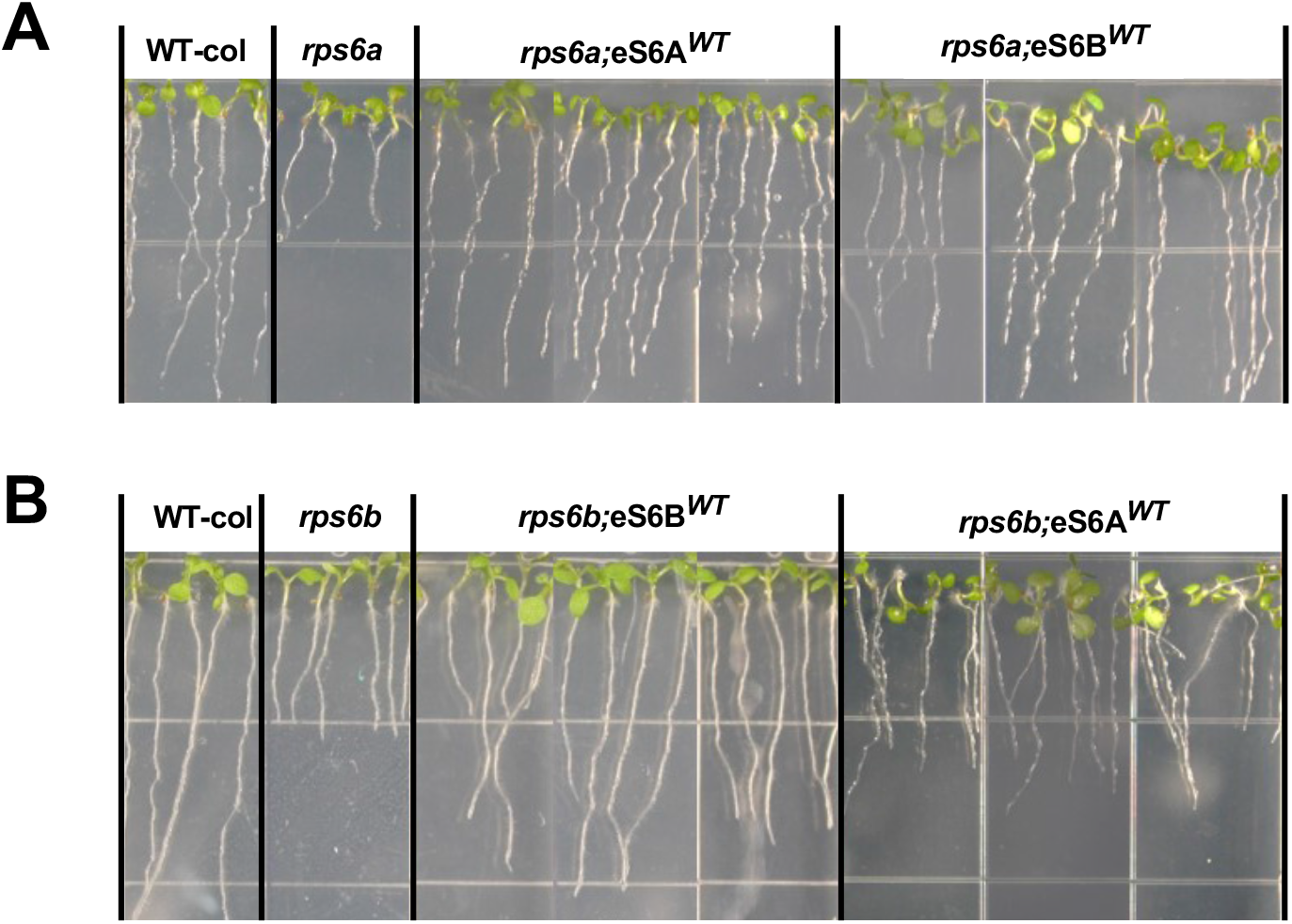
*RPS6A* and *RPS6B* are functionally equivalent. *rps6a-2* (A) and *rps6b-1* (B) mutants were transformed with a transgene of wild-type *RPS6A* or *RPS6B*. Shown are the root lengths of 7-day-old seedlings with their respective wild type (WT) or background mutation.

**Supplemental Figure S4.**
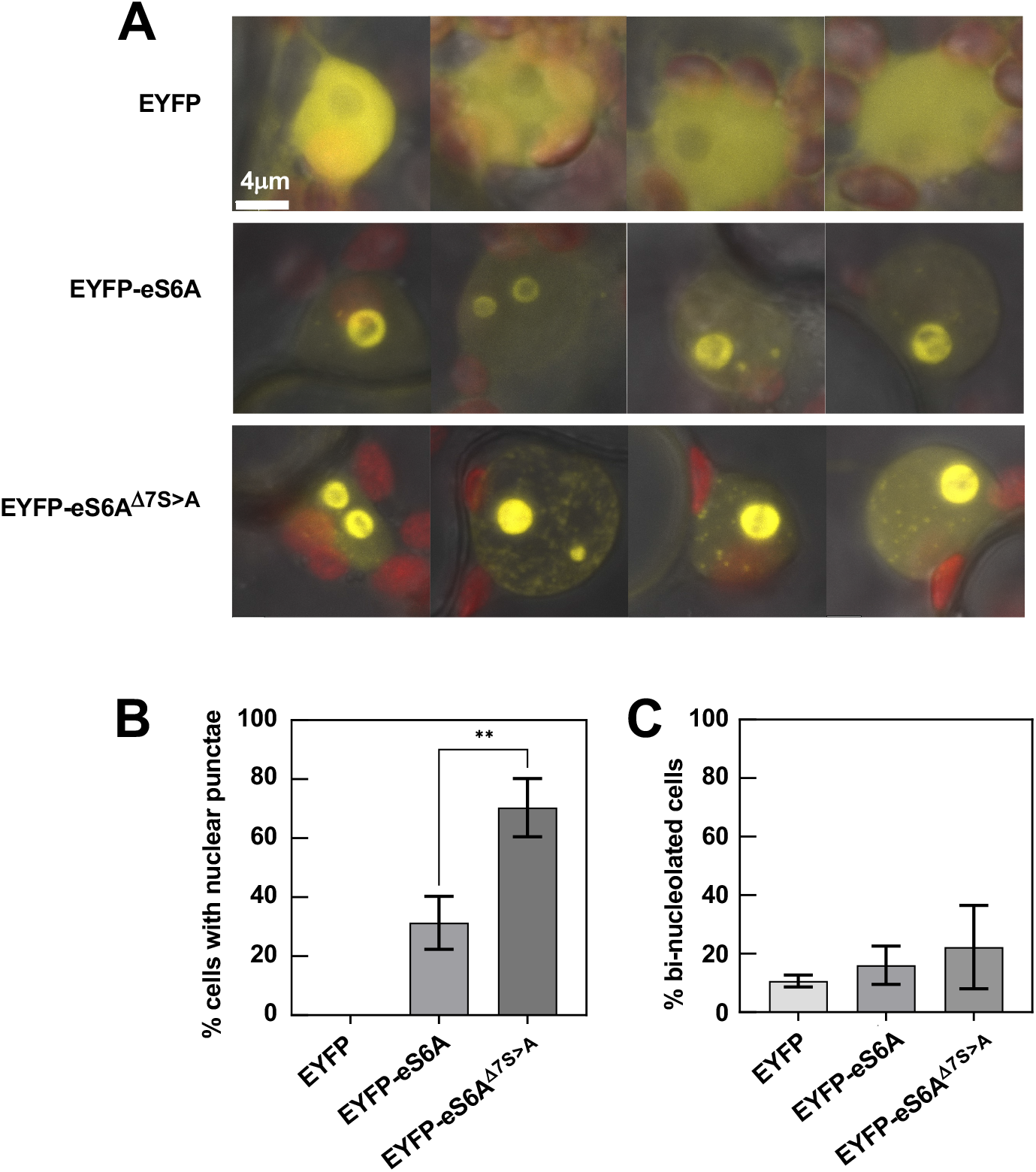
Nuclear and nucleolar localization of YFP-tagged eS6A 7SA in transiently transformed *Nicotiana benthamiana* leaves. Expression was driven by the 35S promoter. Plants were kept in the light prior to microscopic observation to allow for robust phosphorylation of wild-type eS6. Data were collected approximately 48 hours after transformation. Four representative micrographs per construct are shown. Three independent experiments yielded equivalent results. **(A)** The range of typical localization patterns for eYFP alone, wild type eS6A, and eS6A ^1′7SA^. **(B)** Quantification of cells containing nuclear punctae. ** p=0.0073. **(C)** Quantification of the fraction of cells with more than one nucleolus.

**Supplemental Figure S5.**
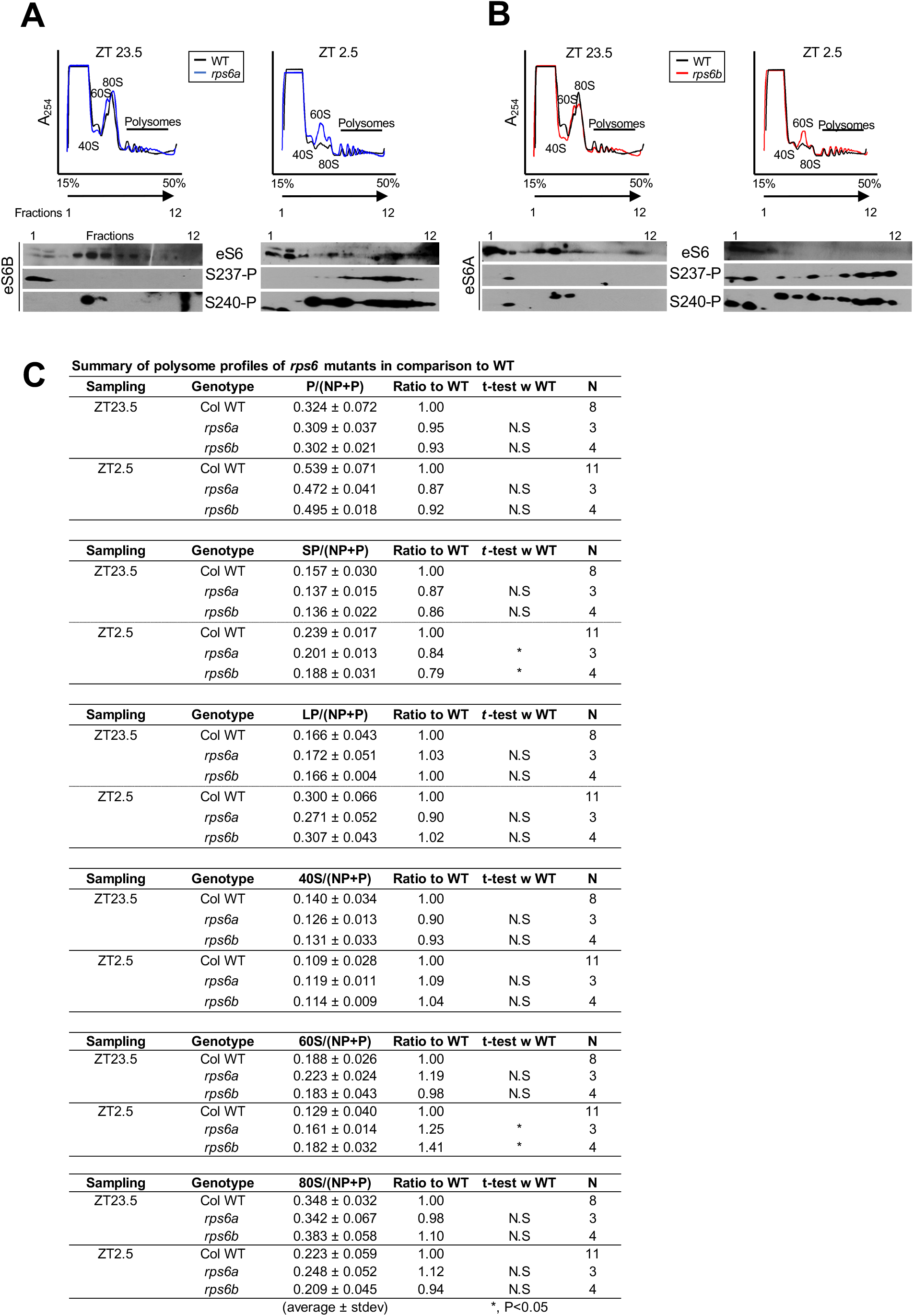
Polyribosome profiles in single *rps6* mutants demonstrate an imbalance in 40S and 60S subunits. **(A)** Ribosome profile of *rps6a* versus wild type at ZT23.5 (30 minutes before lights-on) and at ZT2.5 (150 minutes after lights on). The immunoblots in *rps6a* mutants show the phosphorylation status of eS6B using two phospho-specific antibodies against P-S237 and P-S240 as well as total eS6. **(B)** Same as (A) for *rps6b*. **(C)** Relative abundances of polysomes (P), small polysomes (SP), large polysomes (LP), small ribosomal subunits (40S), large subunits (60S) and entire ribosomes (80S) for WT, *rps6a* and *rps6b*. NP stands for non-polysomal RNA. N stands for the number of replicate gradients; see methods for details.

**Supplemental Figure S6.**
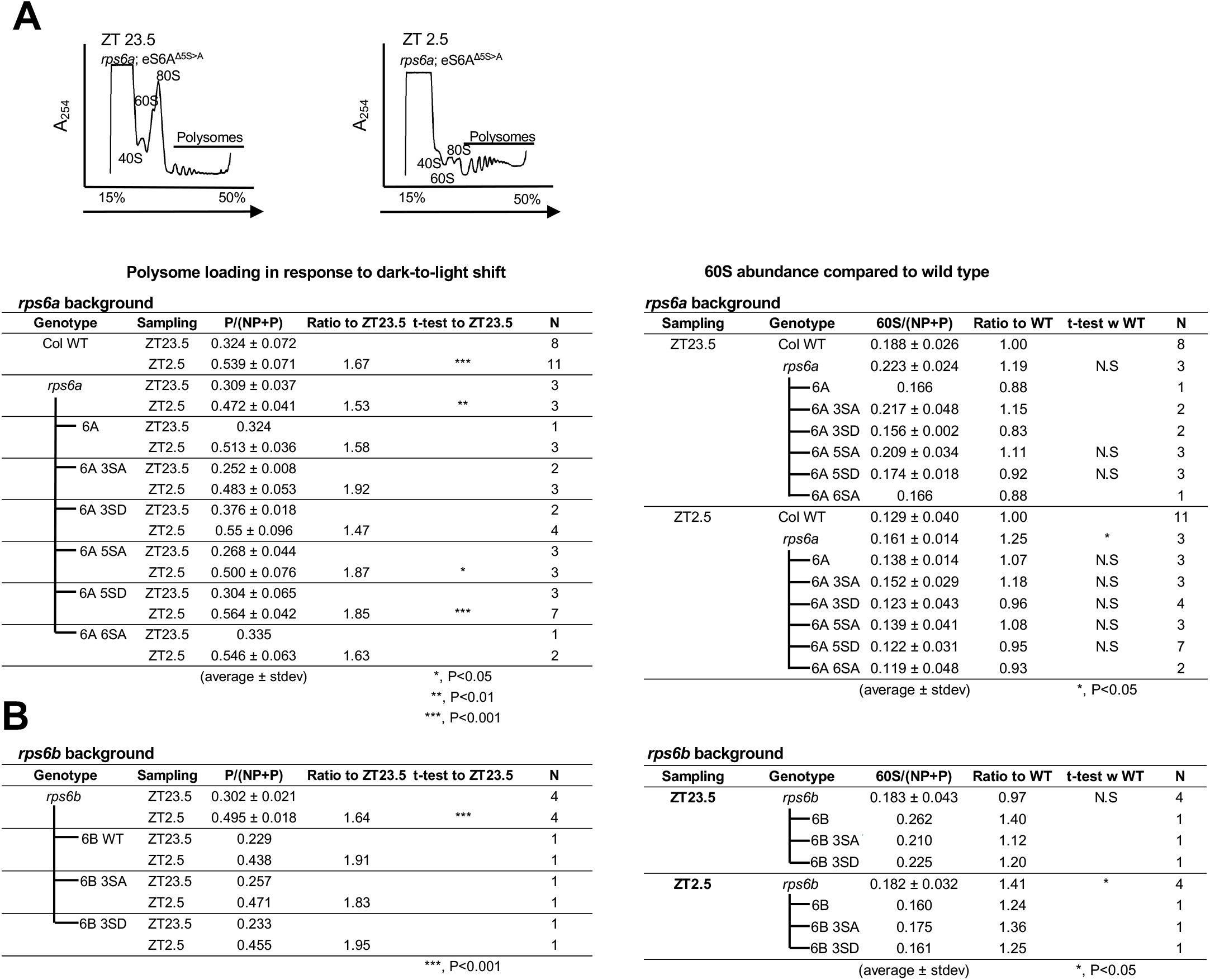
Phosphodeficiency of eS6A has minor effects on global polysome loading. Summary of polysome profiles of single *rps6a* mutants and *rps6b* mutants complemented with various P-enabled and P-deficient eS6 isoforms. N indicates the number of replicate gradients. The left half presents the fraction of RNA in polysomes (polysome loading), focusing on the dark-to-light shift. A representative pair of polysome profiles is shown at the top. The right half quantifies the relative fraction of RNA in the 60S subunit. The data for WT, *rps6a* and *rps6b* are from Supplemental Fig. 5C. **(A)** *rps6a* background. These plants are wild-type for eS6B protein. **(B)** Same as (A) for the *rps6b* background, where the remaining phospho-eS6 is eS6A.

**Supplemental Figure S7.**
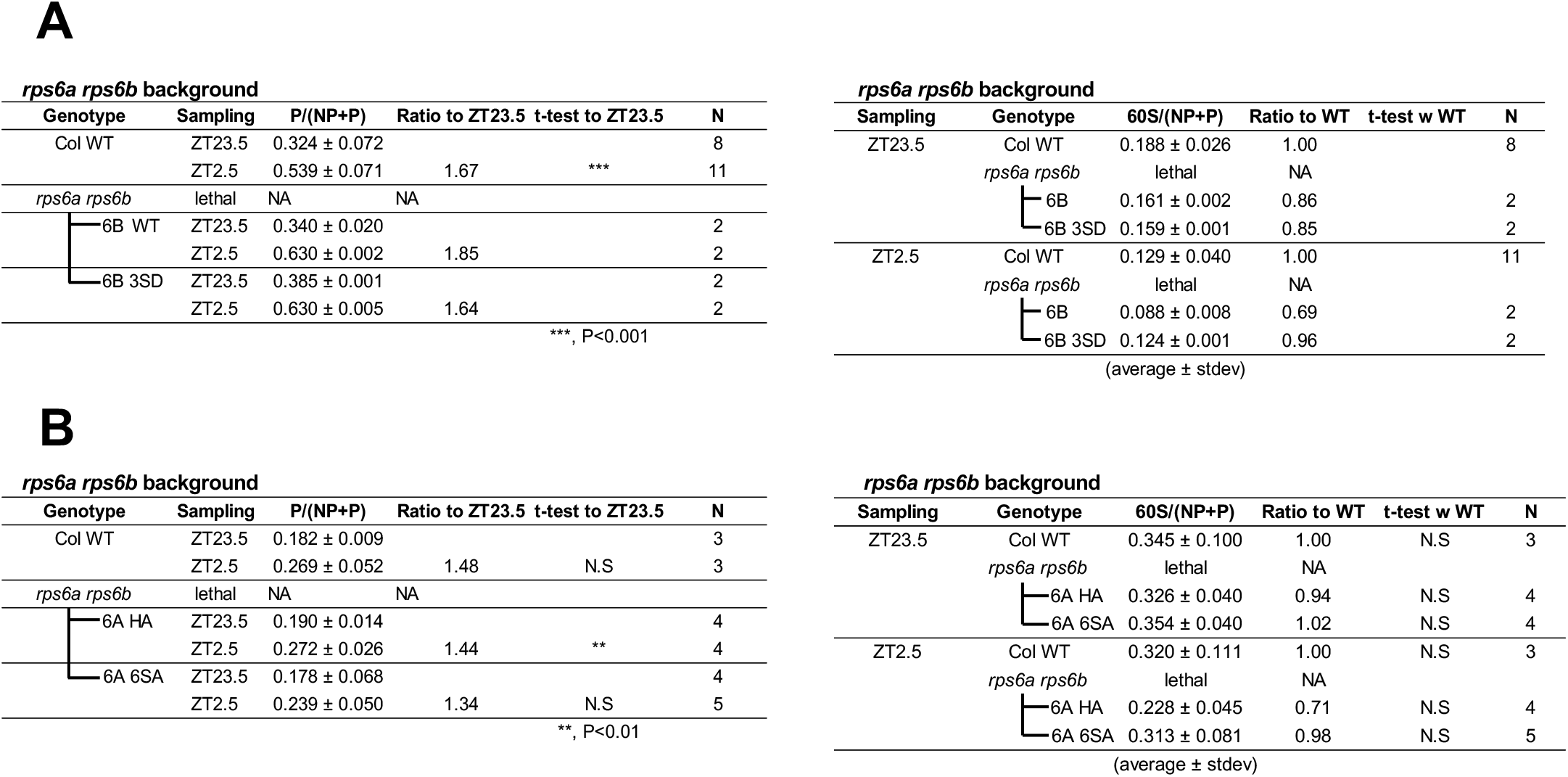
Phosphodeficiency of eS6A has minor effects on global polysome loading. Summary of polysome profiles of *rps6a rps6b* double mutants complemented with various P-enabled and P-deficient eS6 isoforms. N indicates the number of replicate gradients. The left half presents the fraction of RNA in polysomes P/(NP+P), focusing on the dark-to-light shift. The right half quantifies the relative fraction of RNA in the 60S subunit. **(A)** Complementation of double mutants with alleles of eS6B. The data for WT are from Supplemental Fig. 5C. **(B)** Complementation of double mutants with alleles of eS6A. In the 6A 6SA line all eS6 is phosphodeficient.

**Supplemental Figure S8.**
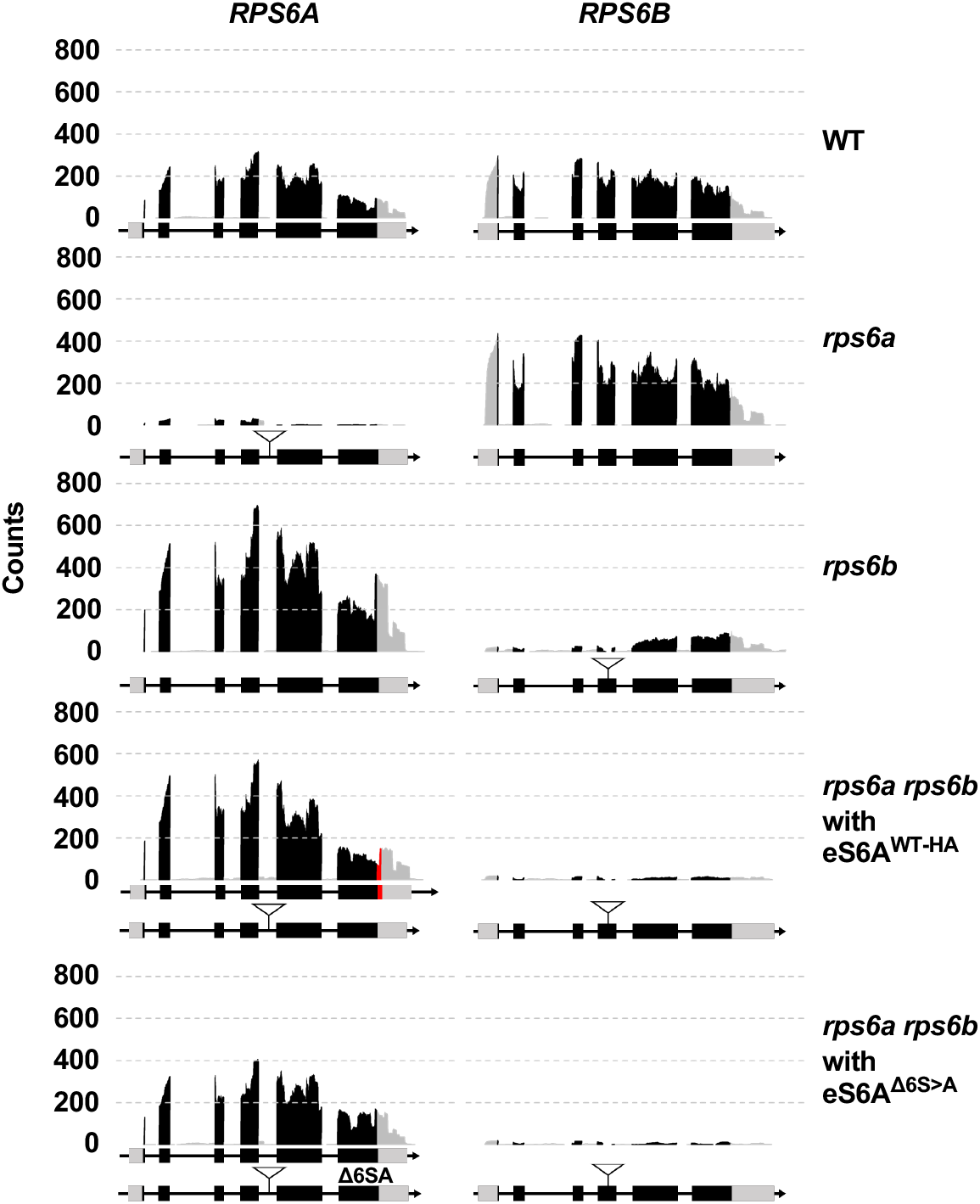
Expression of *RPS6A* and *RPS6B* mRNAs in wild type, the two *rps6* mutants and double mutant plants complemented with eS6^Δ6S>A^ or eS6^WT-HA^. Data are read count pileups from RNA-Sequencing. Note, the residual expression in *rps6b* from the 3’ end of *RPS6B* (see Fig. S1D) was suppressed in the complemented double mutant plants.

**Supplemental Figure S9.**
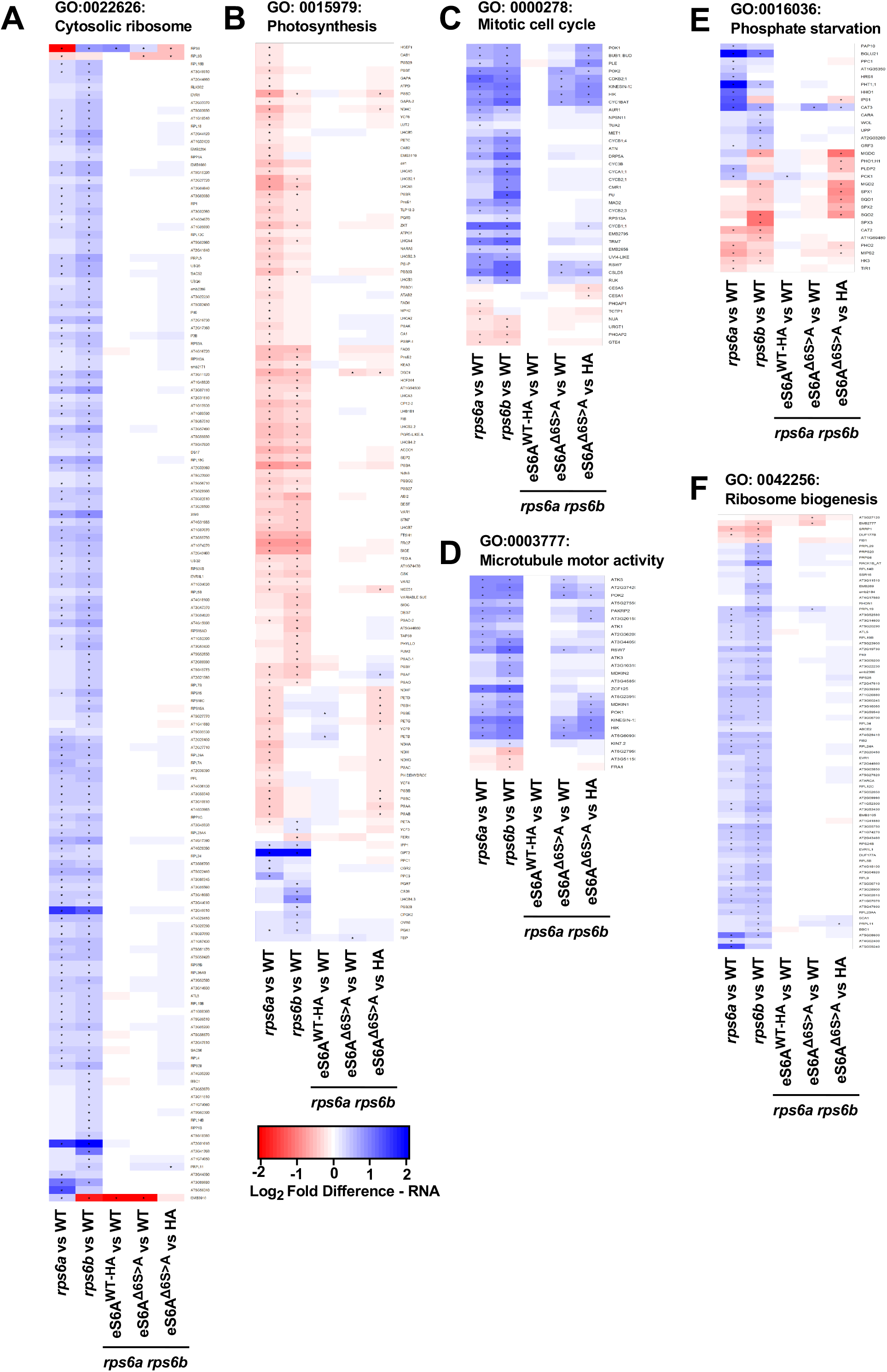
Gene-by-gene heatmaps of differential mRNA expression in eS6 phosphodeficient plants. Genotypes include wild type, rps6a, rps6b and rps6a rps6b double mutants with eS6^Δ6S>A^ or eS6^WT-HA^. Only genes from selected gene ontology terms that passed FDR in at least one of the five pairwise comparisons are displayed. Dots indicate FDR < 0.05. **(A)** Cytosolic ribosome. **(B)** Photosynthesis. **(C)** Mitotic cell cycle. **(D)** Microtubule motor activity. **(E)** Phosphate starvation. **(F)** Ribosome biogenesis.

**Supplemental Figure S10.**
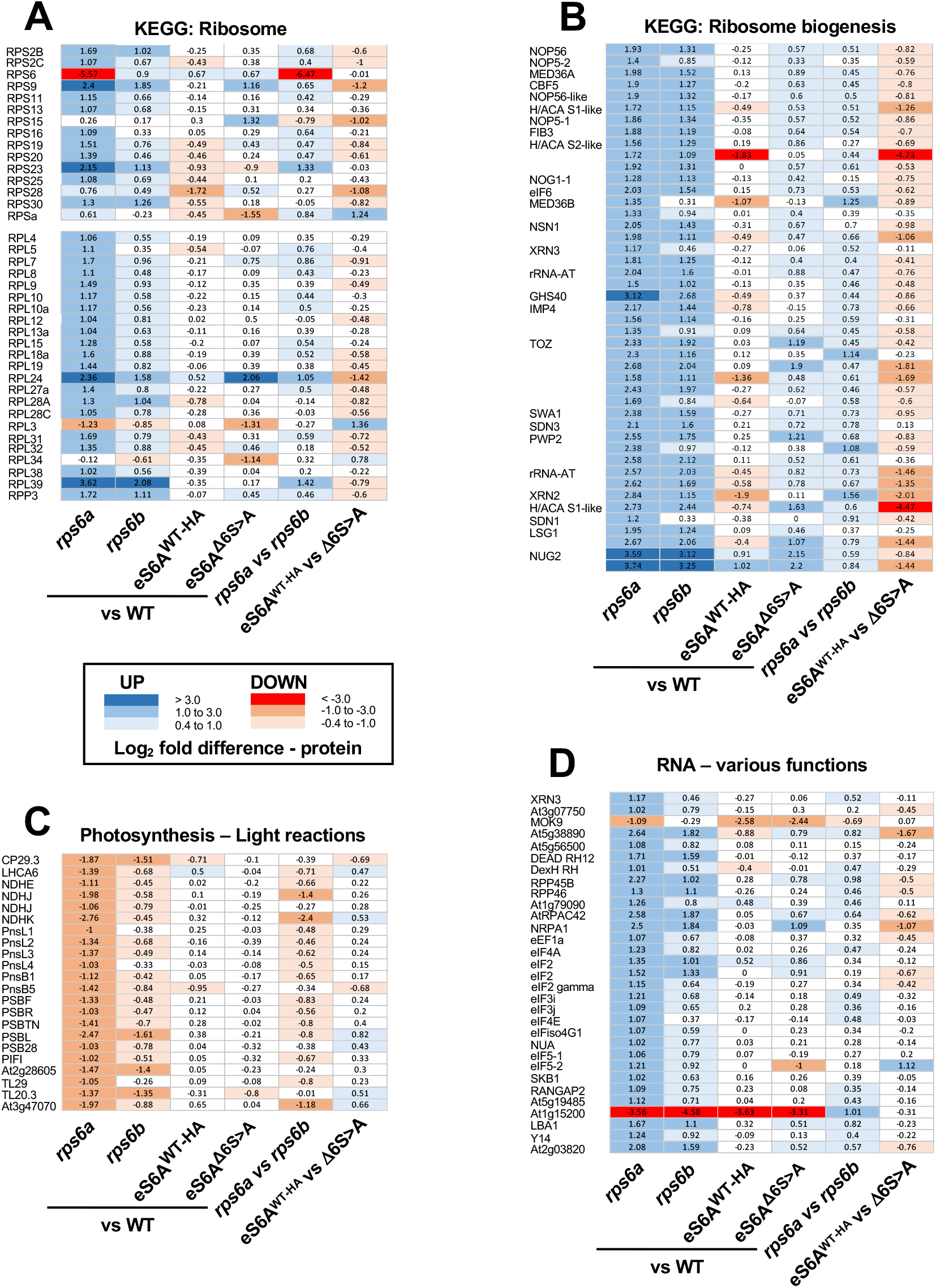
Differential protein abundances detected in 12-day-old seedlings of the indicated five genotypes. Data are log_2_-fold differences to wild type or as indicated. Proteins that are enriched or depleted are shaded blue and red respectively. Selected proteins are annotated with their names. All proteins shown met FDR<0.05 in at least one pairwise comparison.

**Supplemental Dataset 1**. RNA-Seq transcriptome data

**Supplemental Dataset 2**. Differentially expressed proteins

## Supplemental Dataset 1

This table of RNA-Seq transcriptome data displays DESEQ2-normalized read counts for 5 genotypes, WT Columbia, *rps6a*, *rps6b*, *rps6a rps6b* double mutant complemented with a phosphodeficient eS6A^1′6S>A^ allele, and the same double mutant complemented with a phospho-enabled HA-tagged WT allele.

**Supplemental Dataset 2**. Differentially expressed proteins detected by LC-MS/MS in wild type, *rps6a*, *rps6b*, and *rps6a rps6b* eS6A^WT-HA^ and *rps6a rps6b* eS6A^1′6S>A^. All proteins shown met FDR<0.05 in at least one pairwise comparison.

## ACKNOWLEDGEMENTS

We thank the staff of the Oklahoma Medical Research Foundation for RNA-Sequencing services and the Advanced Computing Facility and ISAAC of the University of Tennessee Office of Information Technology for access to high performance computing.

## AUTHOR CONTRIBUTIONS

RE, AD and AGV designed the project. RAUC analyzed data on polysome loading, RNA-Sequencing and proteomics. RE performed research in the early stage of the project. AD performed research in the mid and late stage of the project. SKC contributed the bulk of polysome gradient experiments. LT and JT assisted with experiments under the guidance of AD. PEA performed the proteomics analysis. AGV analyzed data and wrote the manuscript with input from all coauthors.

